# Oatk: a de novo assembly tool for complex plant organelle genomes

**DOI:** 10.1101/2024.10.23.619857

**Authors:** Chenxi Zhou, Max Brown, Mark Blaxter, The Darwin Tree of Life Project Consortium, Shane A. McCarthy, Richard Durbin

## Abstract

Plant organelle genomes, particularly the large mitochondrial genomes with intricate repetitive structures, present significant challenges for assembly. The advent of long-read sequencing technologies provides a transformative opportunity to generate complete genomes, but problems of resolving alternative structures remain. Here we introduce a novel tool for plant organelle genome assembly from high-accuracy long reads. Our method employs a *k*-mer based assembler for rapid assembly graph construction, integrates a profile HMM gene database for robust organelle sequence annotation, and leverages a new search method to find the best supported path through the assembly graph. We describe high-quality organelle assemblies for 195 plant species and demonstrate improvements over other methods. The assembled genomes provide multiple insights into structural complexity, heteroplasmy, and DNA exchange between organelles.

## Introduction

Plastids and mitochondria are integral components of plant cells, governing photosynthesis and respiration, two fundamental physiological processes vital for plant growth and development. They both possess genomes of endosymbiotic origin that have undergone significant reduction and restructuring over evolutionary time ^1^. This includes gene losses, horizontal gene transfers ^1^, intracellular gene transfers ^2^, and inter- and intramolecular recombination ^3^, reflecting a complex evolutionary process shaping their current genomic structure and function. Plastid genomes (plastomes) of land plants are typically 120-160 kb in size and form a highly conserved circular, quadripartite structure, with a large single-copy (LSC) region and a small single-copy (SSC) region separated by two identical (or nearly identical) inverted repeats (IRs) ^4^. In contrast, plant mitochondrial genomes (mitogenomes) demonstrate more extensive size variability, spanning a 200-fold range from tens of kilobases to over 10 megabases ^5^. They can exist in either circular or linear forms, often also contain long exact repeats, and may consist of multiple molecular components, highlighting the diversity and complexity of plant mitogenome architecture ^6^. Beyond their direct importance in plant physiology, organelle genomes provide valuable insights into genetic diversity and evolutionary relationships among plant species. Organelle DNA markers are widely employed in phylogenetic and evolutionary analyses and DNA barcoding initiatives to enhance our understanding of plant biodiversity and taxonomy ^7^.

Organelle genomes have been sequenced using various approaches, starting from Sanger sequencing using primer walking ^8^, to sequencing from purified organelle DNA^9^, and sequencing from whole-cell DNA^7^. While the first two approaches are labour-intensive, costly, and sometimes species-specific, whole genome sequencing (WGS) provides a more efficient, robust and cost-effective way to reconstruct plant organelle genomes. However the long exact repeats in almost all plant organelle genomes, along with other issues discussed below, create severe difficulties for standard genome assembly tools. For this reason there has been a requirement for dedicated software. Most of the contemporary tools for organelle genome assemblies are tailored for WGS data, such as MITObim ^10^, IOGA^11^, NOVOPlasty ^12^, Organelle PBA^13^, and more recently GetOrganelle ^14^, GSAT ^15^, MitoHiFi ^16^ and PMAT ^17^. While the specific implementations may vary, two fundamental components persist in organelle genome assembly from WGS data: (1) distinguishing between organelle-derived and nuclear-derived sequences (potentially from old organelle genome integrations), and (2) assembling reads into genome sequences. The sequence differentiation usually involves the use of a seed database, which may consist of complete organelle reference genomes or sequence fragments derived from conserved genomic regions ^10,11,12,13,14,17^. Raw reads or assembled sequences are classified as organelle sequences if they overlap with these seed sequences. Given the potential divergence between seed sequences and the target genome, an ‘extension’ step is often necessary. This step extends the sequences from the conserved regions with seed hits into neighbouring divergent regions without seed hits by using sequence overlaps. The read assembly often starts with tools originally designed for nuclear genome assemblies, such as SOAPdenovo2^18^ and SPAdes ^19^ used by MITObim, IOGA, GetOrganelle and GSAT for assembling short reads, and the Newbler assembler employed by PMAT for assembling long reads.

Despite the plethora of tools available for organelle genome assemblies, each has inherent limitations. First, most current tools rely on computationally intensive genome assemblers. Although strategies such as assembling only seeded organelle reads or only a limited subset of the WGS data are employed, the computational challenges remain substantial. Second, the ‘seed-extend’ strategy can be problematic, particularly in assembling mitogenomes. Mitogenomes exhibit greater diversity compared to plastomes, posing challenges in creating a universal seed database that accurately represents a wide range of species. Notably, most existing tools, except for GSAT and PMAT, are primarily designed for plastomes and have limited application in mitogenome assembly, tested only on a restricted number of species. Third, many existing tools are optimised for short reads and lack compatibility with long reads, thereby missing a powerful resource for spanning repeats to elucidate complex genome structures. This limitation becomes more pronounced in light of the widespread adoption of highly accurate Pacific Biosciences circular consensus sequence (CCS or HiFi) in genome assemblies ^20,21^. Several tools have explored the use of long reads, but none fully capitalise on the potential of PacBio HiFi reads: Organelle PBA targets long noisy PacBio CLR reads, GSAT utilises long reads only for resolving assembly graphs constructed from short reads, MitoHiFi is only aimed at simple circular mitochondria, and PMAT employs the Newbler assembler, which is slow and less suitable for high coverage data.

Here, we present Oatk to address these challenges. Oatk is a *de novo* organelle genome assembly toolkit for assembling plastid and mitochondrial genomes from WGS data of high-accuracy long reads, specifically the reads used by many recent genome sequencing projects ^22,23,24^. It features a modular design, high speed, and user-friendly interface. Specifically, (1) we have developed a highly efficient genome assembler based on a sparse *k*-mer graph to mitigate the intensive computational demands for read assembly; (2) we have constructed hidden Markov model (HMM) profile gene databases to cover the full breadth of land plant organelle genomes and employ them instead of seed sequences for organelle sequence identification, achieving more accurate classification, particularly for mitogenomes; and (3) we have implemented a sophisticated graph resolution algorithm to generate primary assemblies, taking into account the graph structure and repeat-spanning sequence copy numbers. While the three modules can operate collectively through a wrapper program for organelle genome assemblies, each module can also function independently as a standalone command-line tool, offering enhanced flexibility: the genome assembler can produce assemblies for nuclear genomes comparable to MBG ^25^; the gene database facilitates gene annotation akin to MitoZ ^26^; and the graph resolution module can be utilised in a manner similar to GetOrganelle. We used Oatk to generate organelle genome assemblies for 195 species and compared its performance in read assembly with MBG and PMAT, and in graph resolution with GetOrganelle. Our findings suggest that Oatk generally outperforms the other tools across various metrics. Furthermore, we conducted a survey of the characteristics of the assembled organelle genomes and observed substantial genome diversity both within individual species and between species, for both plastid and mitochondrial genomes.

## Results

### Oatk overview

Oatk consists of three components: (1) *syncasm* for genome assembly using a sparse de Bruijn graph, (2) *hmmannot* for sequence annotation based on a profile HMM gene database, and (3) *pathfinder* for graph resolution leveraging the graph structure and sequence coverage. Below we provide an overview of the key steps and concepts, with details given in the Methods.

For genome assembly, closed syncmers ^27^ with default *k* = 1001, *s* = 31 are first collected from the HiFi sequence data (Fig. 1a) and used to build a sparse de Bruijn graph ^28^ (Fig. 1b). Next, the HiFi reads are mapped to the graph so as to identify low-frequency syncmers derived from sequencing errors and correct them. The graph is then reconstructed from the error-corrected syncmers, followed by graph cleaning and disentangling, and generation of a final assembly graph of unitigs (Fig. 1c). This unitig graph is a general assembly graph including the full nuclear genome. For organelle assembly we next apply a k-mer coverage filter removing unitigs below five times the estimated nuclear haploid coverage depth to remove almost all nuclear sequence, including all but the most recent nuclear mitochondrial or nuclear plastid integration sequences (NUMTs/NUPTs) because *k* is large.

**Figure 1.**
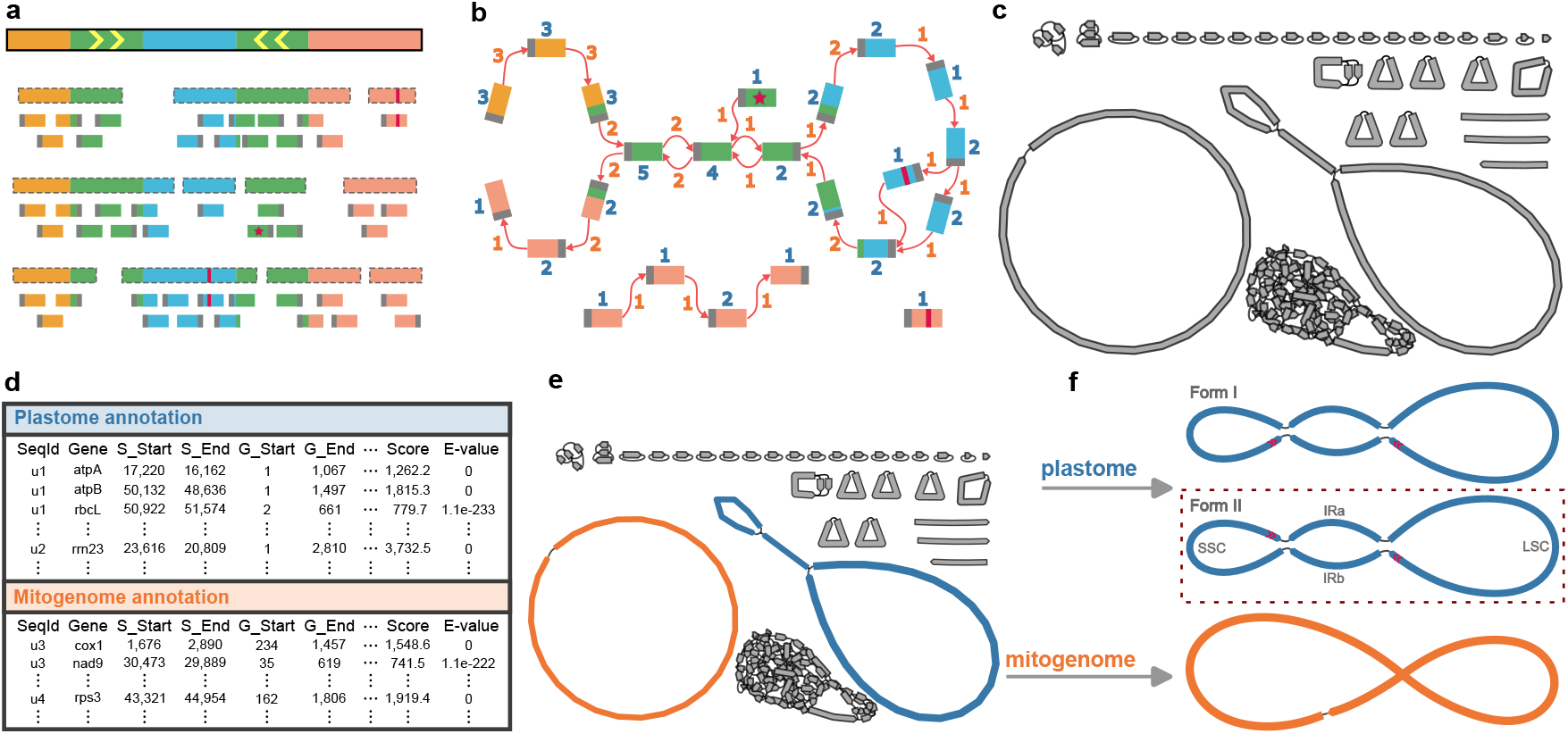
Overview of the Oatk workflow. **a** Collecting closed syncmers from the HiFi sequence data. The top bar with a solid border represents the sequenced genome, with different sequence blocks depicted by colours. The green regions are two IRs, with orientations indicated by the arrows. The bars with dashed grey borders represent HiFi reads, and those without borders represent closed syncmers collected from the reads, with grey solid bars at their ends representing minimal *s*-mers. The red solid bars represent sequencing errors. The syncmer with the red star represents a non-shared terminal syncmer found at the start of a read. Almost all syncmers harbouring sequencing errors or sequence terminals exhibit low frequencies and can be targeted for error-correcting. **b** Constructing a sparse de Bruijn graph from the closed syncmers. Syncmers from (**a**) are connected if they follow in any read. Figures in blue show the copy number of each syncmer, and in orange the copy number of each connection. **c** Assembly graph of high-copy unitigs from moss *Exsertotheca crispa*. In addition to the organelle genomes, a few repeat fragments and ribosomal DNA remain. **d** Generating the gene annotation table by searching the unitig sequences against the gene profile database. **e** Classifying the graph components by organelle types utilising the gene annotations. The blue, orange and grey components are plastid, mitochondrial and non-organelle units. **f** Resolving each organelle graph component to generate individual genome sequences. There are two candidate optimal paths for the plastid graph component, distinguished by the relative orientation of the two single-copy sequences depicted by the red arrows. The gene orders of the two forms are compared to the gene order of the *Arabidopsis thaliana* reference plastome, and the more similar one is selected as the final sequence.

For sequence annotation, each unitig sequence from the assembly graph is searched against a new land plants organelle gene database, comprising 130 plastid and 81 mitochondrial HMM gene profiles (Supplementary Table S1, Supplementary Table S2). The resulting table of gene hits (Fig. 1d) is used to classify each remaining graph component into designated organelle types or non-organelle units (Fig. 1e).

Finally, graph resolution is performed on each organellar graph component to find an optimal path representing a circular or linear form sequence regarded as the primary organelle genome assembly (Fig. 1f). This process considers the graph structure and sequence coverage, involving two major steps: estimating the copy number of each join (edge in the graph) between unitigs (vertices in the graph), and an exhaustive search through graph paths so as to maximise consistency with the edge copy number while covering all sequence. We note that in many cases there is evidence for alternate structures generated by recombination between repeats. We provide a minimal set of best supported structures that represents all sequence in the graph.

### Assembling organelle genomes for 195 plant species

We generated organelle genome assemblies for 195 land plants sequenced by the Tree of Life programme at the Sanger Institute, mostly from the Darwin Tree of Life (DToL) project ^24^, including 24 monocots, 154 eudicots, 16 mosses and one liverwort (Supplementary Table S3). In addition to Oatk, we ran MBG^25^ and PMAT ^17^ for genome assembly construction to compare with the Oatk genome assembler *syncasm*. We also ran GetOrganelle ^14^ for graph resolution to compare with the Oatk graph resolver *pathfinder* (see Methods). The results are summarised in Table 1. Additional details can be found in Supplementary Table S4.

**Table 1.** Organelle genome assembly results of 195 plants.

| Plastome | Oatk | Syncasm | MBG |  | PMAT<sup>1</sup> |  |
| --- | --- | --- | --- | --- | --- | --- |
|  |  | GetOrganelle<sup>2</sup> | Pathfinder<sup>3</sup> | GetOrganelle<sup>2</sup> | Pathfinder<sup>3</sup> | GetOrganelle<sup>2</sup> |
| Single-circular | 192 | 188 | 190 | 185 | 147 | 142 |
| Non-circular | 3 | 6 | 5 | 10 | 16 | 17 |
| Failed<sup>4</sup> | 0 | 1 | 0 | 0 | 32 | 36 |
| **Mitogenome** |  |  |  |  |  |  |
| Single-circular | 132 | 98 | 130 | 83 | 98 | 37 |
| Multi-circular | 29 | 21 | 25 | 17 | 6 | 4 |
| Non-circular<sup>5</sup> | 34 | 67 | 40 | 90 | 59 | 110 |
| Failed<sup>4</sup> | 0 | 9 | 0 | 5 | 32 | 44 |

<sup>1</sup> Only used to generate assembly graphs.
<sup>2</sup> Only used for graph resolution from an input assembly graph.
<sup>3</sup> First ran *hmmannot* to generate annotation results as an input.
<sup>4</sup> Runtime error or reached the 24-hour wall time limit.
<sup>5</sup> Assemblies with multiple components were included if any component was non-circular.

Oatk successfully assembled both plastome and mitogenome for all 195 species. The assembly graphs for these species are presented in Supplementary Figure S1 for plastomes and in Supplementary Figure S2 for mitogenomes. Oatk generated single-circular plastomes for 192 out of the 195 species. The three non-circular genomes belong to *Hibiscus richardsonii, Hibiscus tridactylites*, and *Hibiscus verdcourtii*, the only three Hibiscus species in the dataset. Notably, all three genome assemblies break at the same locus, suggesting either genuine linear structures or issues with the sequencing technique (Supplementary Figure S3). For mitogenomes, Oatk generated 132 single-circular, 29 multi-circular, and 34 non-circular assemblies. Here we classified an assembly as multi-circular if it consists of multiple circular components, and as non-circular if it contains at least one linear component, regardless of whether it has a single component or multiple components.

The results for assembly graph construction using *syncasm* and MBG are comparable. Specifically, the outcomes of the Oatk and the MBG-*pathfinder* combination are similar for both plastome and mitogenome assemblies. GetOrganelle consistently performed better with *syncasm* graphs than with MBG graphs, indicating that *syncasm* outperforms MBG in certain cases. This is anticipated, as *syncasm* has been optimised for organelle genome assembly, particularly for disentangling mixed organelle genome assembly graphs resulting from shared sequences (see Methods, Supplementary Figure S4). While *syncasm* and MBG successfully generated genome assemblies for all species, PMAT failed for 32 species due to either runtime errors or reaching the 24-hour wall time limit. PMAT also shows a high error rate in organelle sequence classification. Out of the 163 species successfully assembled, we observed misclassifications in 147 species (Supplementary Table S5). This is likely because PMAT was initially designed for assembling mitogenomes. We therefore disregarded the classification results of PMAT and used only the assembly graphs as inputs to run *pathfinder* and GetOrganelle. Regarding speed performance, *syncasm* is slightly faster than MBG on average but has higher memory requirements. Both *syncasm* and MBG are significantly faster than PMAT, despite the extensive data downsampling for running PMAT (see Methods, Supplementary Table S4).

For assembly graph resolution, *pathfinder* consistently outperforms GetOrganelle irrespective of organelle types and genome assemblers. While the performance difference is marginal for plastomes, it is notable for more complex mitogenomes. Misidentification of mitochondrial sequences is frequently observed with GetOrganelle, involving both the misidentification of plastid sequences as mitochondrial and mitochondrial sequences as non-organellar (Supplementary Table S4). For instance, GetOrganelle consistently misidentified the IR sequence on the plastomes of moss species as mitochondrial. This is probably because GetOrganelle uses conserved gene sequences as seeds for organelle sequence identification, which may not adequately represent certain species. GetOrganelle was generally faster than *pathfinder*; however the increased time is not a barrier to use of *pathfinder*. In most species, GetOrganelle completed within a few seconds, whereas *pathfinder* took up to a minute (Supplementary Table S4). It should be noted that, for *pathfinder*, the time for running *hmmannot* was included to ensure a fair comparison with GetOrganelle. *Pathfinder* alone typically completed within a second and successfully handled all assembly graphs that GetOrganelle failed to process (Table 1).

### The plastome structures

The majority (182/195) of the assembled plastomes represent a standard quadripartite LSC-IRa-SSC-IRb structure which is clearly revealed by the assembly graphs (Fig. 2a-c). As described in Methods, Oatk outputs these with the standard relative ordering of the large and small single copy regions (LSC and SSC respectively) as in the original *Nicotiniana tabacum* genome ^8^. The other 13 species demonstrate a simple circular assembly graph structure due to the absence of IRs (Fig. 2d), which all fall within the so-called “inverted repeats lacking clade” (IRLC) ^29^. However, since these species are from several genera, their genome sizes vary considerably, ranging from 118 kb in *Erodium maritimum* to 149 kb in *Lathyrus aphaca*. We also observed a broad range of genome sizes for the species with quadripartite structures, from 122 kb in *Lunularia cruciata* to 233 kb in *Schoenoplectus lacustris* (Fig. 2e). Among them, bryophytes have relatively small genomes: the 13 mosses from the class Bryopsida have sizes ranging from 123 kb to 126 kb; the three mosses from the class Sphagnopsida have slightly larger genomes of about 140 kb; the sole liverwort, *Lunularia cruciata*, possesses one of the smallest genomes among all assembled species, at approximately 122 kb, second only to *Erodium maritimum*. In comparison to bryophytes, grasses (Poales) generally have significantly larger genomes, with 13 out of the 19 assembled genomes larger than 180 kb, including 12 sedges (Cyperaceae) and one rush (Juncaceae). The six species smaller than 180 kb are three rushes (*Juncus squarrosus*: 163 kb, *Juncus bufonius*: 170 kb and *Juncus effusus*: 175 kb) and three grasses (Poaceae; *Holcus mollis*: 135 kb, *Bromus sterilis*: 137 kb and *Phragmites australis*: 138 kb). Most of the other species have genome sizes lying between those of bryophytes and Poales, with 136 out of 146 falling within the range of 145 kb to 170 kb, four below this range, and six above it. Specifically, *Calluna vulgaris* and *Jasione montana* have genomes exceeding 200 kb, and mistletoe *Viscum album* represents the smallest genome among all the angiosperms assembled in this study, approximately 129 kb. The expansion of genome size generally resulted in an increased number of protein-coding genes, while the number of unique genes remained relatively stable, with a mean of 80.6 and a standard deviation of 2.0. Bryophyte species, particularly three Sphagnopsida mosses, are exceptions, with 90, 91, and 91 unique genes respectively. In contrast, *Viscum album* lost almost all genes related to NADH dehydrogenase, resulting in a unique gene count of 70, which is well below the average (Fig. 2e, Supplementary Table S6).

**Figure 2.**
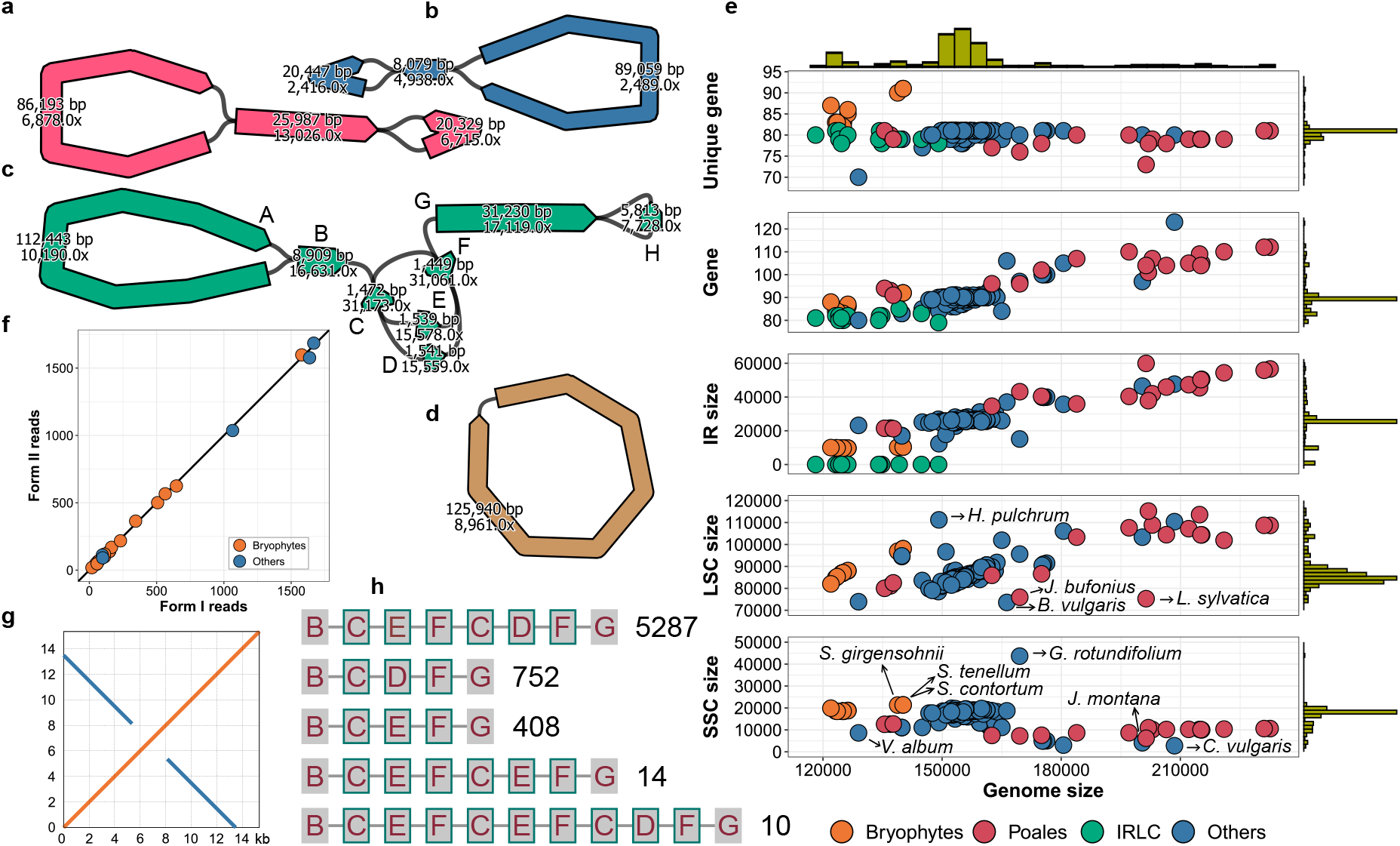
Plastome structures. **a-d** Assembly graphs for four representative structures from the 195 assembled plastomes. The curved bars represent sequences, with the numbers indicating the sequence length and coverage estimated from the syncmer coverage, with the arrow-shaped end indicating the sequence orientation. The thin black lines show the connections between sequences. The assembly graphs were produced using Bandage ^33^ with additional manual adjustments. **a** The *Arabidopsis thaliana* plastome with the regular LSC-IRa-SSC-IRb structure. **b** The *Climacium dendroides* plastome with contracted IRs. **c** The *Calluna vulgaris* plastome with contracted SSC and expanded LSC and IRs, leading to a large genome size of 208 kb. The IR region contains two nested short repeats (C and F) forming a complex structure. **d** The *Medicago arabica* plastome with no IRs. **e** Scatter plots to illustrate the general characteristics of the assembled plastomes as a function of genome size. From top to bottom, the y-axis represents the number of unique protein-coding genes, the total number of protein-coding genes, and the IR, LSC and SSC sizes. The species from the IRLC group are eliminated for the SSC and LSC panels and plotted with value zeros for the IR panel. **f** The number of reads supporting the two differently oriented forms of the plastomes as depicted in Fig. 1f. The 21 species with IRs smaller than 20 kb excluding those from the IRLC group are included in the plot. Each dot represents a species. The black diagonal line indicates identity. **g** The self-alignment of a 15.4 kb HiFi read mapping through the SSC of the *Calluna vulgaris* plastome. The orange and blue lines represent forward and reverse complementary alignments respectively. Referring to **c**, the read maps to the end of IRa (G) for 5.4 kb, then SSC (H) for 2.8 kb, and finally the start of IRb (G from the opposite direction) for 7.2 kb. It should be noted that the SSC size of *Calluna vulgaris* is actually 2,769 bp instead of 5,813 bp as the number depicted in **c** for H. This is because H carries at each end a 1,522 bp sequence belonging to the IRs that forms part of two different syncmers used in the assembly graph construction. **h** Read mapping results in the complex *Calluna vulgaris* IR region grouped by the alignment path. Each block with a label represents a sequence corresponding to those in **c**. Each row represents an alignment path. Each alignment path is followed by a figure indicating the number of reads supporting it. Only reads mapping through this region are counted. Forward mappings (from B to G) and reverse complementary mappings (from G to B) are put into the same group.

Despite having the same quadripartite structure, significant variations in the sizes of different components were observed across species (Fig. 2a-c). Generally, genome sizes positively correlate with the sizes of the LSCs and IRs, but not with the SSCs. This is particularly evident in Poales, where we see notably expanded LSCs and IRs, accompanied by contracted SSCs (Fig. 2e). The average SSC size of the 13 Poales species with genomes exceeding 180 kb is 9.8 kb. In contrast, the average SSC size of the 157 species with genomes smaller than 180 kb, excluding Poales, is 17.9 kb. While this may appear counterintuitive, further investigations suggest that in larger plastomes the SSCs are partially duplicated and transferred into IRs. This IR boundary shift phenomenon is not exclusive to Poales species; it was also observed in other species including *Jasione montana, Calluna vulgaris, Vaccinium vitis-idaea, Inga laurina, Inga leiocalycina*, and *Inga oerstediana*, all of which have plastomes exceeding 175 kb. Moreover, the shifting of IR boundaries in these species is asymmetrical with respect to the two IR/SSC boundaries. The boundary adjacent to the *ndhF* gene remains relatively stable, with duplication always occurring from the opposite end of the SSC (Supplementary Figure S5). The proportion of the SSC that was duplicated varies across species. In extreme cases, the SSC contracted to such an extent that only the *ndhF* gene remains, as observed in *Jasione montana* (4.1 kb), *Vaccinium vitis-idaea* (3.1 kb), and *Calluna vulgaris* (2.8 kb, Fig. 2c and Fig. 2g). The dynamics of plastomes extend beyond IR/SSC boundary shifts, encompassing a broad spectrum of phenomena. For example, in *Berberis vulgaris*, the IRs shift into the LSC, while in *Luzula sylvatica*, the IRs shift into both the LSC and SSC. In *Hypericum pulchrum* and *Geranium rotundifolium*, the LSC and SSC shift into the IRs, accompanied by intricate genome rearrangements, resulting in anomalous component sizes. In *Viscum album*, the SSC is highly contracted, leading to the loss of all genes related to NADH dehydrogenase (Fig. 2e, Supplementary Figure S6-Supplementary Figure S10).

In addition to the structural dynamics observed across different species, we also detected abundant heteroplasmy within individual species. From the point of view of the assembly graph, the single copy regions SSC and LSC can in principle be traversed in two distinct relative orientations, indicating that the plastome graph represents two distinct structural configurations (Fig. 1f). It has long been reported that both forms can coexist within a single individual, a phenomenon that could be explained by flip-flop recombination between two IRs ^30,31^. Recently, Wang and Lanfear ^32^ revisited this discovery using a method combining long-read mapping and statistical modelling and found that this type of heteroplasmy exists in most plants. Here we utilised assembly graphs to facilitate a more direct investigation of this topic. Since a read must map through the entire IR region in the graph to provide evidence for a specific form, we set the IR size limit to 20 kb taking into account the length restriction of PacBio HiFi reads, resulting in a selection of 23 species, including 18 bryophytes and five angiosperms. For each species, we mapped the HiFi reads to the individual assembly graph and count the number of reads supporting each form. For all species tested we see approximately the same number of reads supporting both forms (Fig. 2f) with no significant deviation from a binomial distribution with probability 0.5 (Supplementary Table S7). This observation reaffirms previous findings that the two forms widely coexist in plants, with their ratio being close to 50/50 and is consistent with frequent intramolecular flip-flop recombination as a plausible explanation for this phenomenon. Besides chromosomal-scale structural variations, we also observed other types of heteroplasmy at smaller scales. Figure 2c shows the assembly graph of *Calluna vulgaris* as an example. The IR region contains two nested small repeats, leading to a complex structure. By mapping the reads to this region, we identified multiple isoforms with varying molecule abundances (Fig. 2h). Similar heteroplasmic structures were observed in several other assembled species (Supplementary Figure S1, Supplementary Figure S11).

### The mitogenome structures

The assembled mitogenomes are much more variable than the plastomes, demonstrating remarkable diversity in size, structure, and gene content (Fig. 3a-g, Supplementary Figure S2). Genome sizes range from 103 kb to 2,731 kb, with the smallest genome observed in the moss *Thuidium tamariscinum* and the largest in the sedge *Carex laevigata*. A similar trend in genome sizes was observed in mitogenomes as in plastomes, with bryophytes having relatively small mitogenomes, Poales mitogenomes being notably larger on average, and other species falling between them. There is a positive correlation between gene number and mitogenome size in general (Fig. 3g, Supplementary Table S8). Among the 195 mitogenomes analysed, 132 represent a single circular structure. These single circular mitogenomes exhibit various underlying structures, including simple ring structures, quadripartite structures with either an inverted or directed repeat (DR), and more complex structures involving multiple circular molecules linked by repeat sequences (Fig. 3a-d). Twenty-nine of the assembled mitogenomes consist of multiple circular components (Fig. 3e). Notably, the *Galeopsis tetrahit* mitogenome exemplifies this complexity, comprising 368.8 kb of sequence distributed across ten circular molecules, each containing protein-coding genes (Supplementary Figure S12). The remaining 34 assembled mitogenomes are non-circular, comprising either a single linear component or multiple components, with at least one being linear. While some of these non-circular mitogenomes are probably the consequence of incomplete assembly, others can be verified through read mapping. For instance, the *Alnus glutinosa* mitogenome consists of two components: one circular and one linear, with the latter displaying an intriguing loop structure at each end (Fig. 3f). Reads were found mapping through each direction of the bifurcation in the loop structure, with a smooth change in coverage, confirming the authenticity of this structure (Fig. 3h). The loop structure is not unique to *Alnus glutinosa*; similar structures were observed in at least six other species assembled in this study (Supplementary Figure S2).

**Figure 3.**
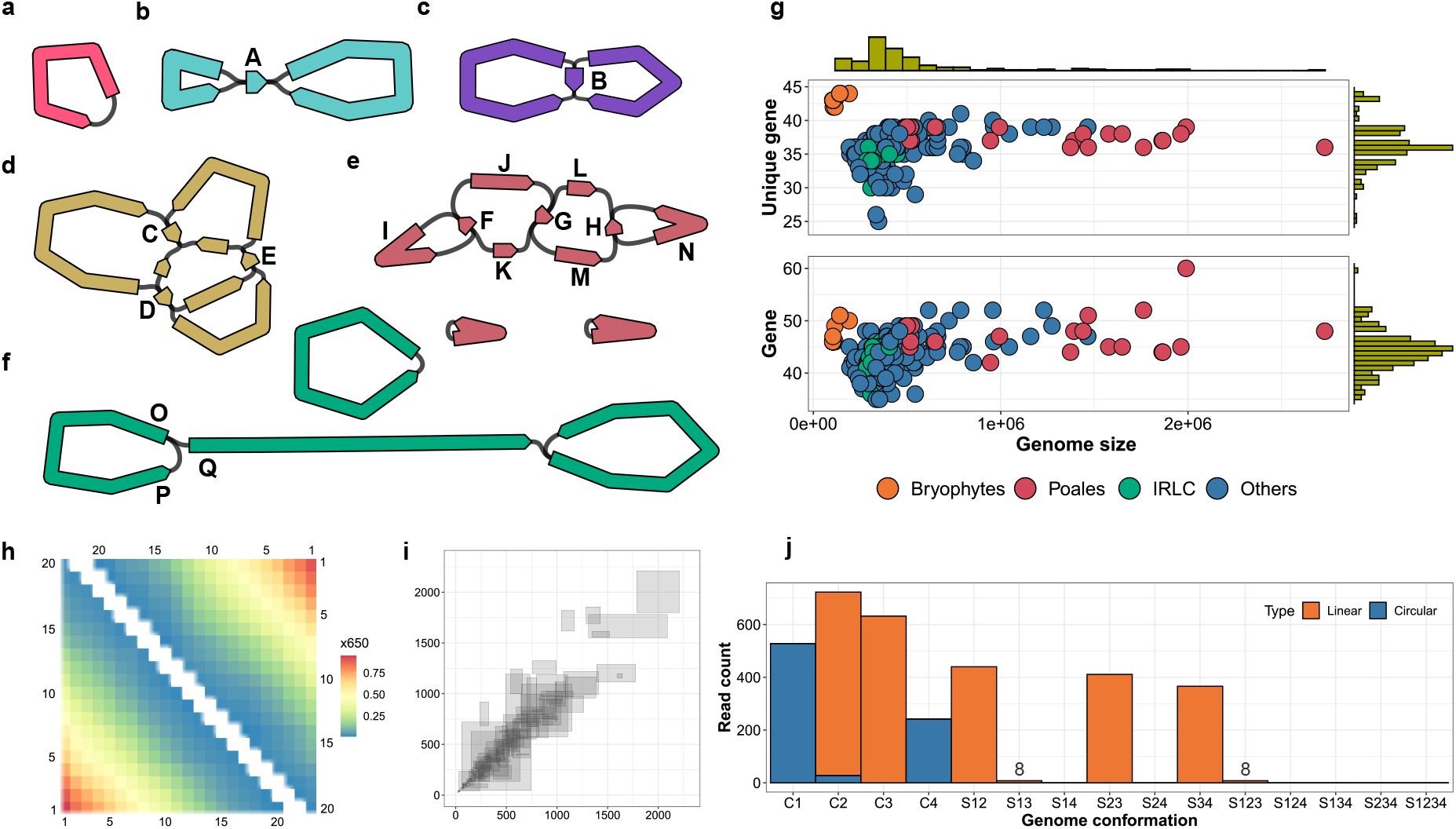
Mitogenome structures. **a-f** Assembly graphs for five representative structures from the 195 assembled mitogenomes. The assembly graphs were produced and notated as described in Fig. 2. **a** The *Climacium dendroides* mitogenome with a single circular structure. **b** The *Pseudognaphalium luteoalbum* mitogenome with a quadripartite structure featuring an IR. **c** The *Calluna vulgaris* mitocgenome with a quadripartite structure featuring a direct repeat. **d** The *Quercus robur* mitogenome with a complex structure featuring multiple repeats. The genome can be resolved into a single circular structure. **e** The *Erodium maritimum* mitogenome with three components that all can be resolved into circular structures. **f** The *Alnus glutinosa* mitogenome with two components that can be resolved into a circular and a linear structure. **g** Scatter plots to illustrate the unique protein-coding gene number (top panel) and the total protein-coding gene number (bottom panel) in the assembled mitogenomes as a function of genome size. **h** A heat map to illustrate the results of read mapping across the graph junctions O-P (lower triangular) and O-Q (upper triangular) as depicted in **f**. The axes represent the distances to the junction point up to 20 kb, divided into 20 intervals, each spanning 1 kb. Each cell denotes the count of reads covering the respective intervals in two sequences. For instance, the cell at coordinates (3,5) indicates the number of reads spanning the O-P junction and covering both the 3 kb interval on sequence O and 5 kb interval on sequence P. **i** A scatter plot to illustrate the read mapping results on the 165 repeats identified in the assembled genomes that bifurcate bidirectionally. Sequences A to H in panels **b** to **e** exemplify such repeats. It should be noted that the sequences connected to the repeat may originate from the same sequence approached from two directions, as seen in cases A, B, F and H. Let **R** denote the repeat sequence, **IA, IB** two incoming sequences, and **OA, OB** two outgoing sequences. Each grey-shaded rectangle in the plot corresponds to a repeat sequence, with left, right, bottom and top boundaries indicating the number of reads supporting **IA**-**R**-**OA, IB**-**R**-**OB, IA**-**R**-**OB** and **IB**-**R**-**OA**, respectively. **j** A bar chart to illustrate the read mapping results supporting structural heteroplasmy for the graph depicted in **e**. The x-axis categorizes different heteroplasmic subgraph structures: C1 to C4 correspond to four minicircles featured respectively by non-repeat sequences I, J and K, L and M, and N; ‘S’ categories represent supercircles formed by multiple minicircles. For instance, S24 denotes the double-circle structure comprising C2 and C4. A read is considered to support a minicircle if its mapping path covers at least two non-repeat sequences, including covering the same sequence twice (e.g., I-F-I for C1). A read is considered to support a supercircle if it covers at least one non-repeat sequence from each constituent minicircle. A read mapping is classified as linear if it starts and ends with different sequences, and circular if with the same sequence.

Structural heteroplasmy was commonly observed in the assembled mitogenomes. The genome of *Pseudognaphalium luteoalbum* exhibits a standard plastid-like quadripartite structure with IRs (Fig. 3b). Similar to observations in the plastome, this structure manifests in two coexisting forms, characterised by different orientations of the two single-copy sequences. The number of reads mapping across the IRs supporting the two forms are 299 and 301, respectively, suggesting unbiased interchange. We observed a similar phenomenon in the genome of *Calluna vulgaris*, which also represents a quadripartite structure but with DRs (Fig. 3b). The number of reads supporting the two forms are 328 and 361, respectively. Unlike the IR-featured structure, in this scenario, the second form consists of two separate small circles, supported by 170 and 191 reads, respectively. We extended this analysis to all accessible bidirectionally bifurcated repeats across the assembled mitogenomes. Specifically, we examined each sequence on the assembly graphs and included it as a repeat in the analysis set if it meets the following criteria: (1) the sequence is no larger than 10 kb; (2) the sequence has two incoming edges and two outgoing edges; and (3) the size of each sequence associated with the four edges is at least 10 kb. The sequence size threshold was set to ensure the quality of read mapping. This process identified 165 repeats across 109 genomes. The read mapping results suggest that nearly all these repeats are associated with structural heteroplasmy, with many representing nearly balanced abundances (Fig. 3i, Supplementary Table S9). The repeat-driven dynamics of local structures can extend globally, leading to diverse molecular forms involving various combinations of subgenomic regions. (Fig. 3j). The coexistence of these superstructures and subgenomic structures has been noted previously, with one possible explanation being intramolecular recombination ^6^.

### Sequence transfer between organelle genomes

We observed frequent sequence sharing between plastid and mitochondrial genomes within a species, reflecting historical transfer of DNA from one organelle to the other ^34^. This usually led to a mixture of sequences from two organelle genomes in the same assembly graph forming a entangled structure. Figure 4a demonstrates the genome assembly graph of the *Alnus glutinosa* as an example. It contains four identical shared sequences with sizes from 1,382 bp to 4,753 bp. The homologous sequence pair forms a bubble structure, differentiated by a few variants, including seven SNPs and three 4-7 bp small INDELs. For each species, we aligned the plastid and mitochondrial genomes to identify shared sequences. We set a minimum sequence size of 1 kb and a minimum sequence identity of 90% to eliminate noisy alignments. We identified 576 shared sequences totaling 1.57 Mb across 144 species, with individual shared sequence sizes ranging from 1 kb to 17.3 kb. Among them, 57 species contains more than 10 kb of shared sequences, with *Carex laevigata* having the most at 66.6 kb. This finding demonstrates the prevalence of sequence transfer between organelle genomes, although, notably, no shared sequences are found in any of the bryophyte species. No clear correlation between mitogenome size and the amount of shared sequences was observed (Fig. 4b). For example, *Erodium maritimum* has a 307.7 kb mitogenome and 37.9 kb of shared sequences, whereas *Juncus bufonius* has a 993.7 kb mitogenome but only 3.4 kb of shared sequences. The majority of sequence transfers occurred from plastome into mitogenome, not the other way around. Out of the 1.57 Mb of shared sequences identified, 1.10 Mb were annotated as gene sequences, with 1.09 Mb (98.7%) specifically identified as plastid genes (Supplementary Table S10).

**Figure 4.**
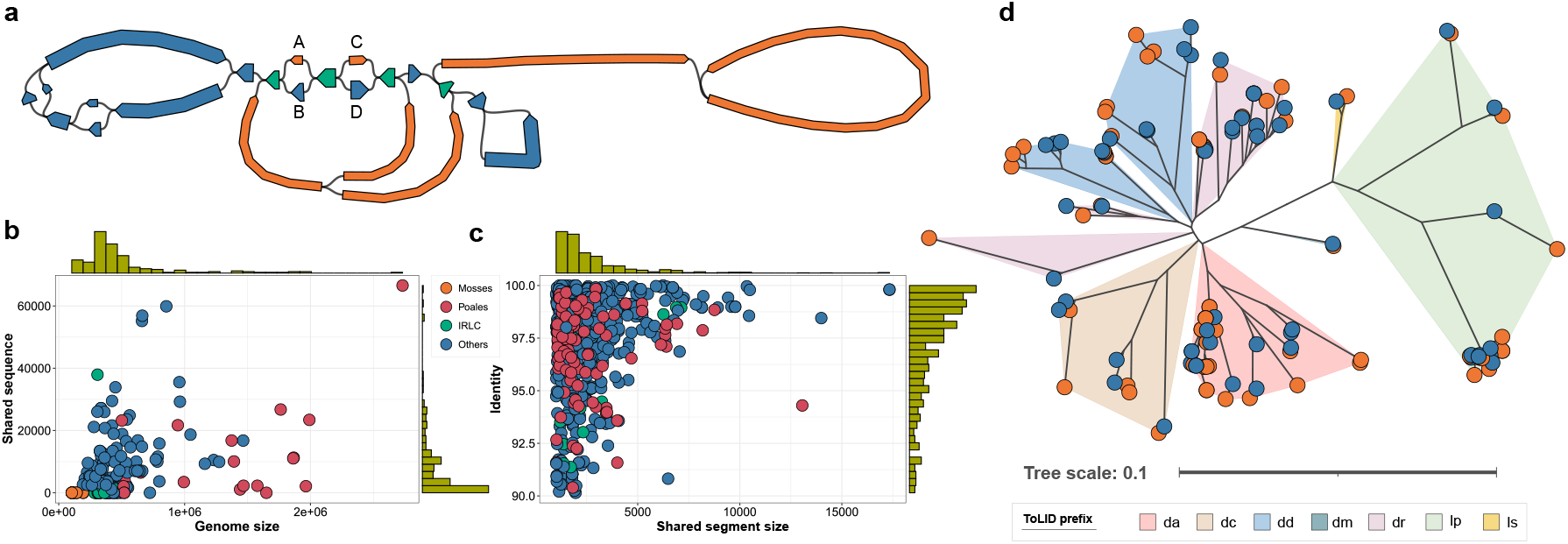
Sequence transfer between organelle genomes. **a** The *Alnus glutinosa* organelle genome assembly graph before disentangling. The circular component of the mitogenome is omitted from the plot for clarity (Fig. 3f). Plastid sequences are coloured in blue, mitochondrial sequences in orange, and shared sequences in green. The assembly graph was produced and notated as described in Fig. 2. **b** A scatter plot to illustrate the total size of the shared sequences between the plastid and mitochondrial genomes for each species as a function of the mitogenome size. **c** A scatter plot to illustrate the sequence identity for each homologous sequence pair as a function of the size of the shared sequence segment. **d** A phylogenetic tree to illustrate the relationship among 128 homologous *psaA* gene sequences. Each leaf node denotes a sequence, coloured blue or orange to indicate plastid or mitochondrial genomes, respectively. The ToLID group prefix (https://id.tol.sanger.ac.uk) is employed to delineate clades. The tree plot was produced using iTOL^35^ with additional manual adjustments.

In terms of sequence similarity between paralogs, 287 sequences totalling 895.5 kb show greater than 98% identity, while 161 sequences totaling 223.2 kb show less than 95% identity. Although many paralogs represent high similarities, sequence identities are broadly distributed across the entire range from 90% to 100%, indicating that sequence transfer is a repeated, ongoing process (Fig. 4c). This is supported by phylogenetic analysis of 64 paralogous sequence pairs annotated as the *psaA* gene - the most frequently annotated shared gene followed by *psbB* with 58 sequence pairs. The phylogenetic tree constructed from these 128 sequences accurately represents the clade structure and shows a stratification of shared sequences (Fig. 4d). For many paralog pairs, two sequences were closely clustered together in the tree. Notably, sequences in 19 paralog pairs are mutually the nearest neighbour of each other and sequences in 33 paralog pairs are mutually within the three-nearest neighbours (Supplementary Table S11). Clades for some genera like *Hypericum* and *Inga* show separate clusters for plastid and mitochondrial sequences, suggesting that sequence transfer occurred before their speciation. In 50 out of the 64 paralog pairs, the mitochondrial sequences demonstrate greater distance to the node representing the most recent common ancestor (MRCA) compared to their plastid paralogs, such as observed in species *Juncus squarrosus, Chamaenerion angustifolium*, and *Iris foetidissima* (Supplementary Table S12). Statistical analysis using the Wilcoxon signed-rank test on all 64 paralog pairs indicates a significant difference in distances (*p*-value = 8.969e-06). This suggests that those sequences transferred into mitogenomes evolve faster than their counterparts in plastomes.

## Discussion

We developed Oatk for *de novo* assembly of plant organelle genomes using high-accuracy long reads and applied it to assemble 195 species spanning a broad range across the tree of life. Compared to other tools, Oatk produced complete assemblies for more species, particularly for mitogenomes. Oatk is distinguished from other tools by a highly efficient genome assembler utilising a sparse *k*-mer graph to reduce memory usage and accelerate graph construction. For *k*-mer selection, we use closed syncmers, which are intrinsically suitable for *k*-mer sparsification ^27^. The sparsification factor is approximately (*k* − *s* + 1)*/*2, where *k* and *s* are the *k*-mer and *s*-mer size respectively. With high-accuracy long reads, we can choose a large *k* and typically have *k* ≫ *s*. In this case, the average distance between adjacent *k*-mers is roughly *k/*2. For example, with *k* = 1, 001 as used in this study, the data is compressed by a factor of about 500, thereby facilitating efficient assembly graph construction. Other techniques for *k*-mer sparsfication include the minimizer approach used in MBG^25^ and the *k*-min-mer approach used in MDBG ^28^. MBG features a syncmer-based implementation when integrated into Verkko ^36^. The major difference between *syncasm* and the syncmer-based MBG probably lies in graph disentangling, with *syncasm* being better tailored for organelle genome assemblies.

Even though there are many well-designed genome assembly tools for HiFi data, such as hifiasm ^37^ and HiCanu ^38^, they have not been effectively used for organelle genome assemblies. These assemblers are not optimized for extremely high coverage data, such as organelle genomes, where coverage can reach thousands or tens of thousands of times, leading to problems with sequence error correction. Additionally, the highly unbalanced data coverage between organelle genomes and nuclear genomes presents another challenge. Finally, these tools generally rely on string graphs, which suffer from inaccurate sequence coverage estimation - a critical factor for graph resolution in plant organelle genome assemblies. In contrast, *k*-mer based assembly graphs provide better sequence coverage estimates.

Another distinctive characteristic of Oatk is its use of a profile HMM gene database for organelle sequence identification, rather than relying on seed sequences. For a given gene, a profile HMM was constructed considering all related sequences available in the NCBI repository to ensure a broad species representation. This approach provides a more sensitive method for sequence identification, particularly useful for divergent mitogenomes. Oatk is not the first tool to use profile HMMs for genome annotations. They have been employed by MitoZ to identify target sequences in animal mitogenome assembly ^26^, and by GeSeq as an essential component for protein and rRNA coding gene annotation in plant organelle genomes ^39^. Although our focus has been on plant organelle genome assemblies, Oatk extends its utility to other species with the corresponding gene database. We have created HMM profile gene databases for various clades and developed a tool to facilitate the creation of gene databases given an NCBI taxonomy ID (Methods). As an example, we applied Oatk to assemble complete mitogenomes for five animal species, including a mammal, a fish, a bird, a lizard and an insect (Supplementary Figure S13). This demonstrates the broader applicability of Oatk beyond plants.

We observed extensive diversity in the assembled genomes, both within and across species. Intriguingly, analyses combining long reads and assembly graphs offered significant insights into the structural heteroplasmy of these genomes, highlighting their potential as powerful tools for understanding complex organelle genome structure. However, this poses an important question regarding the appropriate representation of plant organelle genomes. Traditionally, genomes are stored as linear sequences and labelled as circular if applicable, as required by public sequence repositories in the International Nucleotide Sequence Database Consortium (INSDC: NCBI GenBank, ENA and DDBJ), whereas the structural heteroplasmy information present in organelle genomes is overlooked. Some tools, such as GetOrganelle and GSAT, opt to output sequences of all possible conformations ^14,15^, a practice Oatk could potentially adopt. However, this approach introduces additional challenges such as sequence redundancy and combinatorial complexity, especially in the case of complex mitogenomes where the number of conformations may escalate to an unmanageable level. We suggest that it might be more useful to provide a primary assembly that includes all sequences together with the assembly graph, allowing flexibility depending on the research question. This underscores the need for tools that can directly analyse assembly graphs, such as for gene annotation, where considering graph structure may change annotation outcomes.

## Methods

### Collecting closed syncmers and representing reads by syncmer vectors

Closed syncmers ^27^ are parameterised by two positive integers *s* and *k* (*s < k*) and a hash function *φ* of strings of length *s* that returns the same value for a string and its reverse complement. Given a string *κ* of length *k*, also called a *k*-mer, to determine if *κ* is a closed syncmer, *φ* is first used to compute the hashes for all *s*-long substrings (*s*-mers) from *κ*. The *k*-mer *κ* is a closed syncmer if and only if the hash of the first or last *s*-mer of *κ* is minimal, i.e. it is not larger than all other hashes from *κ*. We use *k* = 1, 001 and *s* = 31 in this study. Given a set *R* of HiFi sequencing reads, we build a *k*-mer table *T* containing all the distinct closed syncmers present at least once in the reads. Since most HiFi sequencing errors are homopolymer run length errors ^20^, the reads are homopolymer compressed (hoco) before collecting syncmers, where homopolymer runs are collapsed into a single nucleotide. In *T*, each *k*-mer is assigned a unique sequential index. The table also maintains the the *k*-mer frequency (coverage), its hoco sequence, and the its full length consensus sequence. To generate the consensus sequence of a hoco *k*-mer, the homopolymer run lengths for each base of the *k*-mer are recorded during the *k*-mer table construction. The run length consensus of a base is computed as the closest integer to the average of all run lengths at its respective positions, mirroring the approach used in MBG ^25^. We use a hash function, denoted by *ψ*, to map *k*-mer sequences to 64-bit integers for faster comparisons during the table construction. It is similar to *φ* for *s*-mers, excepting that while *φ* is designed to be a perfect hash function eliminating hash collisions, hash collisions for *ψ* are inevitable, owing to the values of *s* and *k*. The hash collisions are checked, and distinct table entries are created for *k*-mers with collided hashes. To allow for both directions of double-stranded DNA, *ψ* is applied to compute hash values for both the original *k*-mer and its reverse complement, and only the sequence with the smaller value constitutes a *k*-mer entry in the table.

Given the *k*-mer table *T*, each read in the read set *R* can be rewritten as a vector of syncmers accompanied by the respective positions on the read and orientations concerning the DNA strands. Let *r* ∈ *R* be a HiFi read harbouring *m* closed syncmers, denoted by *κ*_1_, *κ*_2_, …, *κ*_*m*_ sequentially. The respective positions of these syncmers on *r* are *p*_1_, *p*_2_, …, *p*_*m*_ in ascending order. Let *π* be a function mapping a *k*-mer to its index in the table. Let *ρ* be a function calculating the relative orientation of a *k*-mer compared to the corresponding entry in the table, with 0 being the same strand and 1 being the reverse complement. The read *r* can then be rewritten as *ζ* (*r, T*) = [⟨*π*(*κ*_1_), *p*_1_, *ρ*(*κ*_1_)⟩, ⟨*π*(*κ*_2_), *p*_2_, *ρ*(*κ*_2_)⟩, …, ⟨*π*(*κ*_*m*_), *p*_*m*_, *ρ*(*κ*_*m*_)⟩], where *ζ* is the transformation function. Let *Z* = *ζ* (*R, T*) be the transformed read set after applying *ζ* to each read in *R*. The inherent characteristic of assured overlap between consecutive closed syncmers ^27^, i.e., *p*_*i*+1_ − *p*_*i*_ *< k* for all *i* ∈ [1, *m*), guarantees that the whole read apart from material before the first syncmer and after the last syncmer can be restored from its respective syncmer vector.

### Constructing a sparse *k*-mer graph from closed syncmers

The *k*-mer table *T* and the transformed read set *Z* are used to construct a sparse *k*-mer graph. For each *k*-mer entry *κ* in the table *T*, two vertices are established, one for the *k*-mer itself and the other for its reverse complement 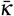 denote these as *v* and 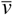 respectively as an example. Vertices *v* and 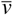 are called complement vertices to each other, and 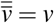 holds. Vertices maintain references to the *k*-mer indices, allowing for retrieval of *k*-mer information from the table, including details such as the sequence and the coverage. Two complement vertices refer to the same *k*-mer entry in the table. However, it is necessary to calculate the reverse complements when retrieving the underlying *k*-mer sequence and *k*-mer consensus sequence for the one not present in the table.

Directed edges are introduced between vertex pairs when the corresponding *k*-mers are adjacent on a transformed read. For any edge, a complement edge is also added. Consider the example of the read *ζ* (*r, T*). Let *v* and *w* be the associated vertices of the first two *k*-mers ⟨*π*(*κ*_1_), *p*_1_, *ρ*(*κ*_1_)⟩ and ⟨*π*(*κ*_2_), *p*_2_, *ρ*(*κ*_2_)⟩, respectively. Given the read *r*, a directed edge *v* → *w* will be added to the graph, pointing from vertex *v* to *w*, and simultaneously its complement edge 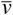 is added, where 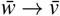 is the respective complement vertex of *v*. Edges record their coverage representing the count of positions on the transformed reads supporting the adjacency of the two vertices. In the example, the read *ζ* (*r, T*) contributes coverage of one unit to both the edges *v* → *w* and 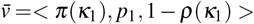.In addition, edges retain the length of the sequence overlap between the *k*-mers represented by the two vertices. For a given edge, the sequence overlap is determined by consolidating all overlap distances calculated from the positions of the two corresponding adjacent *k*-mers on the transformed reads. Typically, all the overlaps for a particular *k*-mer pair edge will be identical, and this uniform distance will be utilised to calculate the overlap length. However, in rare cases, it is possible to observe multiple overlap lengths due to complex repeat structures; when this happens the largest distance will be employed to calculate the overlap length. The complement edges *v* → *w* and 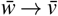 exhibit identical edge coverage and sequence overlap length by definition.

Let *G* = (*V, E*) be the *k*-mer graph, where *V* is the vertex set and *E* is the edge set. The above construction procedure guarantees *G* is Watson-Crick complete ^40^: (i) 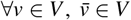;and (ii) 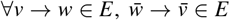;where 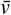 and 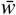 are the complement vertices of *v* and *w*, respectively. For all *v* ∈ *V*, let *δ* ^+^(*v*) be the outdegree of *v* and *δ* ^−^(*v*) be the indegree. It follows that 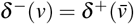.

### Correcting syncmer errors

The *k*-mers in the table *T* with coverage below a predefined threshold *ϑ* are identified as potential error syncmers. Let Θ be the set of potential error syncmers. Firstly, the vertices corresponding to the *k*-mers in Θ are deleted in the *k*-mer graph *G*. Denote by *G*^′^ the trimmed *k*-mer graph, assumed to be free of syncmer errors. Then, for each read in *Z*, error blocks, defined as consecutive blocks of error syncmers, are identified and mapped to *G*^′^ for error correction. Consider the example of the read *ζ* (*r, T*), let *β* = [*κ*_*a*+1_, *κ*_*a*+2_, …, *κ*_*a*+*b*_] be an error block of *b* syncmers, where 0 ≤ *a < a* + *b* ≤ *m*. By definition of the error block, (i) ∀*i* ∈ [1, *b*], *κ*_*a*+*i*_ ∈ Θ; (ii) *κ*_*a*_ ∉ Θ if *a >* 0; and (iii) *κ*_*a*+*b*+1_ ∈*/* Θ if *a* + *b < m*. For simplicity, we have omitted the mapping functions *π* and *ρ* here for converting actual *k*-mers to *k*-mer table entries. We call *κ*_*a*_ the left boundary of the error block and *κ*_*a*+*b*+1_ the right boundary. If both the left and right boundaries are absent, the entire read constitutes an error block and will be excluded from error correction. Without loss of generality, we assume the existence of the left boundary; otherwise, we can establish the left boundary by converting the read to its reverse complement. The syncmer vector *κ*_*a*_ ∪ *β* delineates a subsequence of the read *r* harbouring sequencing errors, starting from the position *p*_*a*_ and concluding at the position *p*_*a*+*b*_ + *k*. Let *ξ* be the subsequence, and *l* = |*ξ* | = *p*_*a*+*b*_ − *p*_*a*_ + *k* be the length in base pairs of the subsequence. In broad terms, the error correction aims to find an unambiguous syncmer path over *G*^′^ starting with the vertex corresponding to the left boundary *κ*_*a*_ outlining a sequence with a sufficiently small Levenshtein edit distance to *ξ*. If the right boundary *κ*_*a*+*b*+1_ exists, the syncmer path must end with the vertex corresponding to it. The upper limit of the Levenshtein distance is parameterised by *ε* and calculated as *εl* for a subsequence of length *l*. The parameter *ε* should reflect the overall base accuracy of the sequencing data. In this study, we set *ϑ* = 3 and *ε* = 0.01.

To find the error-correcting syncmer path for an error block, we perform a depth-first search (DFS) on *G*^′^ starting from the vertex corresponding to the left boundary, denoted by *v*_0_. Let *v*_0_, *v*_1_, …, *v*_*n*_ be the current search path of the DFS at depth *n*. Let *ξ*_*n*_ be the corresponding sequence delineated by the search path, and *l*_*n*_ = |*ξ*_*n*_| be the sequence length. A dynamic programming (DP) table is first constructed for calculating the Levenshtein distance between *ξ* and *ξ*_*n*_. Let *d*_*n*_ = *η*(*ξ, ξ*_*n*_) be the edit distance of the prefix alignment between two sequences, i.e., the edit distance ignoring the trailing indels in the DP table. Let *g*_*n*_ = *γ*(*ξ, ξ*_*n*_) be the length of the overhang sequence of *ξ*, i.e., the number of trailing insertions of *ξ* in the DP table. The actual edit distance between *ξ* and *ξ*_*n*_ is computed as *e*_*n*_ = *d*_*n*_ + *g*_*n*_. The ‘depth search’ along the path is concluded at *v*_*n*_ under the following conditions: (i) *d*_*n*_ ≥ *εl*, or (ii) *l*_*n*_ ≥ (1 + *ε*)*l*, or (iii) there is no succeeding vertex from *v*_*n*_. When a ‘depth search’ is completed, the subpaths along the search path are examined to update the global minimum and the second minimum edit distances, denoted by *ê* and 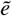,respectively. The path with the minimum edit distance is also updated. The DFS search routine then proceeds to the next search path. A constraint on the total number of ‘depth searches’, parameterised by *τ*, is set to prevent excessive path exploration in complex graph regions arising from highly repetitive sequences. The error correction is considered unsuccessful if the number of searched paths exceeds the constraint. Upon the successful completion of the DFS, the values of *ê* and 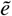 are examined. The error correction is considered valid if *ê* ≤ *εl* and 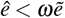,where *ω* is a parameter to control path ambiguities, with lower values indicating higher confidence of the optimum path. For a valid error correction, the path with the minimum edit distance is used to update the syncmer vector of the read. In this study, we set *τ* = 10, 000 and *ω* = 0.7. The bottleneck in error correction arises from the computation of Levenshtein distance. We implemented two strategies to address this issue. Firstly, we employ Ukkonen’s *O*(*ND*) algorithm ^41^, where *N* is the minimum length of two sequences and *D* is the edit distance between them. The fundamental idea involves constraining the DP matrix computation to a band along the diagonal that is 2*D* wide. This implementation is intrinsically suitable for our needs, considering we set an upper limit of *εl* for the Levenshtein distance given a sequence of length *l*. Secondly, we reuse the DP matrix during the DFS. When visiting a vertex at each DFS step, the target sequence is extended by appending the sequence encoded in the current vertex, and the computation of the DP matrix is confined to only the part relevant to the newly added bases.

### Cleaning a k-mer graph and generating a unitig graph

Given that organelle genomes typically exhibit significantly greater coverage than the nuclear genome in the sequence data, we set a lower threshold for *k*-mer coverage, denoted by *c*, to exclude *k*-mers derived from the nuclear genome. After filtering the low-copy *k*-mers, we apply a strategy similar to Miniasm ^40^ to remove tipping sequences shorter than ten kilobase pairs and pop bubbles shorter than 100 kilobase pairs. A unitig is defined by a ‘maximal unambiguous path’ in the *k*-mer graph. More precisely, a vertex path *v*_0_ → *v*_1_ → … → *v*_*n*_ forms a unitig if *δ* ^+^(*v*_*i*_) = *δ* ^−^(*v*_*i*+1_) = 1 for ∀*i* ∈ [0, *n*) and (i) *v*_0_ = *v*_*n*_ or (ii) *δ* ^−^(*v*_0_) ≠ 1 and *δ* ^+^(*v*_*n*_) ≠ 1. Since each vertex in the *k*-mer graph corresponds to a *k*-mer entry in the *k*-mer table, a unitig can be written as a syncmer vector, similar to a read. Let *U* be the set of unitigs. For ∀*u* ∈ *U*, let *ζ* (*u, T*) be the syncmer vector for *u*, using the same notations for reads. A unitig graph is derived using *U* as vertices, and for ∀*u*_0_, *u*_1_ ∈ *U*, adding a directed edge *u*_0_ → *u*_1_ pointing from *u*_0_ to *u*_1_ if the last *k*-mer of *u*_0_ is identical to the first *k*-mer of *u*_1_. The structure and characteristics of the *k*-mer graph persist in the unitig graph. Specifically, the unitig graph remains Watson-Crick complete. To generate the sequence for a unitig, the *k*-mer sequences along the *k*-mer vector are concatenated, eliminating the overlapping sequences between each pair of adjacent *k*-mers. The final genome assembly comprises the sequences of all unitigs. For a complement unitig pair, only one copy is retained in the assembly to avoid redundancy.

### Resolving complex assembly structures

Horizontal transfers of DNA between organellar genomes can result in an entangled assembly graph for the two organelle genomes owing to the presence of shared sequences longer than the *k*-mer used for graph construction. To tackle this problem, we leverage the phase information of *k*-mers obtained from reads represented by vectors of *k*-mers. Firstly, the reads are mapped to the unitig graph at the k-mer level employing a seed-and-extension approach: the first *k*-mer of the read presented in the unitig graph is used as the seed to initiate a DFS; the DFS path is then extended based on the order of *k*-mers on the read. Then, for each unitig node, the number of reads supporting each spanning triplet are counted and used to resolve the local graph structure around the node. A spanning triplet of a node (the centre node) is a graph path consisting of three nodes with an incoming and an outgoing node connecting the centre node.

For example, node *u* has *m* = *δ* ^−^(*u*) incoming nodes, written as 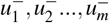, and *n* = *δ* ^+^(*u*) outgoing nodes, written as 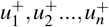,then there are *m* × *n* spanning triplets for *u*. Write 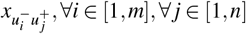, as the number of reads supporting the triplet path 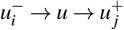.The triplet path 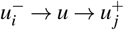 is called dominated if 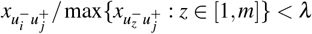 and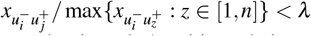, where 0 *< λ* ≤ 1 is a predefined threshold and set as 0.1 in this study; otherwise it is non-dominated. A unitig node is called resolvable if there exists at least one dominated spanning triplet. If a node is resolvable, we update the unitig graph by introducing a compound node for each non-dominated triplet. Precisely, for a non-dominated triplet 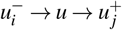,the compound node, denoted by *υ*, is constructed by concatenating the *k*-mer vectors of 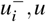 and 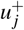 i.e.,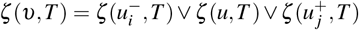 all the incoming edges pointing to 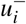 will be copied and point to *υ*; all the outgoing edges pointing from 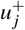 will be copied and point from *υ*. The resolved node *u* is finally removed from the unitig graph. This resolution process is repeated for multiple rounds until the graph structure converges.

### Constructing a profile HMM gene database

We construct the profile HMM gene databases separately for plastid and mitochondrial genomes. Given the species and organelle genome type, we download all related sequences from the NCBI repository in Genbank format using the Entrez Direct command tools ^42^. Specifically, we use the *esearch* tool with the query string “TAXID [Organism] AND OTYPE [Filter]” to search against the nucleotide database for a list of target sequences and the *efetch* tool to download those sequences, with ‘TAXID’ being ‘txid3193’ for Embryophyta, ‘txid40674’ for Mammalia, ‘txid7898’ for Actinopterygii, ‘txid8782’ for Aves, ‘txid8504’ for Lepidosauria and ‘txid50557’ for Insecta, and with ‘OTYPE’ being ‘chloroplast’ or ‘mitochondrion’. A sequence set is created for each gene, including protein-coding genes, tRNA genes, and rRNA genes, after parsing the Genbank file and used to construct a profile HMM. For profile HMM construction, we first perform an initial sequence clean by removing sequences with invalid characters and those shorter than one-third or longer than three times the average length. If the number of sequences is more than 10,000 after the initial clean, a second-round clean is performed to retain the top 10,000 most representative sequences, determined by 12-mer completeness: a 12-mer table is first generated for all sequences, excluding the 1% most and 1% least frequent ones; a score is then computed for each sequence as the sum of counts for all unique 12-mers; the 10,000 sequences with the highest scores are finally chosen. With the cleaned sequences, the multiple sequence alignment is constructed using MAFFT ^43^ (version 7.505) with the ‘--auto’ option and the profile HMM is built using HMMER’s ‘hmmbuild’ tool ^44^ (version 3.4) with default parameters. All gene profile HMMs are concatenated into a single file and compressed with HMMER’s ‘hmmpress’ tool (version 3.4) to make a binary database.

### Annotating organelle genome sequences with the gene database

The assembled sequences are searched against the profile HMM database with HMMER’s ‘nhmmscan’ tool (version 3.4) to build a table for protein-coding genes. Gene hits with *e*-values greater than 1 × 10^−6^ or scores less than 300 are considered insignificant and omitted in the further analysis. The gene annotations are used to determine the organelle type for each assembly graph component. We construct two gene tables for each graph component, one for plastid genes and one for mitochondrial genes, containing only the best hit of each gene on the sequences belonging to the component. We could simply assign the organelle type of a component by comparing the sum of scores to plastid and mitochondrial genes. However, this sometimes leads to misassignments of mitochondrial sequences into plastid due to the frequent insertion of plastid sequences into mitochondrial DNA. Since plastid genes are well conserved, they have higher annotation scores than mitochondrial genes on average. As a result, a few inserted plastid genes can become dominant. To mitigate this problem, we use a progressive assignment approach rather than a simple comparison of the sum of scores. The graph components are sorted by sum of scores in descending order and processed progressively: (1) if the component has not been processed, assign it the corresponding organelle type; (2) if the component has been processed and classified as plastid, reassign it to mitochondrial if (i) there has already been a component assigned to plastid, and (ii) the mitochondrial score is no smaller than one-third of the plastid score. The above process could incorrectly identify some junk sequences as organelle sequences, including those originating from NUMT/NUPT or resulting from sequencing errors. To address this issue, we track the highest score for each gene in sequences already assigned to an organelle type. When we need to assign an organelle type to a new component, we compare the gene scores in sequences from this component to the current highest scores. A component is assigned only if it has a sufficient number of gene hits with higher scores than the current best. We set this threshold at three for plastid and one for mitochondrion. The highest scores for the genes are updated each time a new component is added.

### Constructing a primary sequence assembly from the unitig graph

The genomes of plant organelles frequently cannot be assembled into a single contiguous sequence because of large repeats or alternative isoforms. We examine all possible path conformations within the assembly graph, selecting the longest path as the primary genome assembly. If multiple paths are of equal length, we choose the one most strongly supported by the read data, determined by the total number of reads spanning the node junctions along the path. We start by estimating the copy number of each sequence in the graph. Consider an assembly graph of *n* nodes, denoted by *u*_1_,…,*u*_*n*_. The length of *u*_*z*_ is *l*_*z*_ and the sequence depth of coverage of *u*_*z*_ is l*d*_*z*_, *z* = l 1, …, *n*. We estimate the average base sequence coverage, denoted by 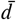, and determine the copy number of *u*_*z*_ as 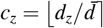. This estimation is performed using an EM algorithm. In the E-Step, we calculate 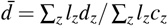.and in the M-Step, we calculate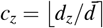. With the copy number information for each sequence, we aim to traverse the entire graph while ensuring each node is visited according to its copy number. To achieve this, we expand the graph by creating additional copies of multi-copy nodes and their corresponding edge connections. In the expanded graph, each node only needs to be visited once. We implemented the path exploration algorithm to ensure that: (1) all paths are examined; (2) equivalent paths due to rotation and reversal are examined only once; and (3) equivalent paths resulting from permutations of different copies of the same nodes are examined only once. During the path exploration, the circular and linear paths encompassing the longest sequences are added to the candidate set for the ultimate selection. For the final choice of the optimal path, a circular path is preferable over a linear path when the circular one covers no less than a predefined portion (90% in this study) of the sequences covered by the linear one in length; among the paths of the same length, the path with the most read support (designated as the graph edge coverage) is preferable over the others.

In plastome assemblies, there are typically two possible path conformations due to IRs. We select the one that shows better gene order consistency with the *Arabidopsis thaliana* reference plastome ^45^, which has the same relative order as the original published *Nicotiniana tabacum* plastome ^8^. Specifically, we use the order of the 71 protein-coding genes from the *A. thaliana* assembly as a reference. We compute Spearman’s rank correlation coefficients between the reference and each of the path conformations and select the one with the higher correlation coefficient. Additionally, we rotate the assembled sequence so that it begins with the start codon of the *psbA* gene as a convention.

If the primary sequence assembly resulting from the above process is non-circular, we adjust the sequence copy number estimates using the graph topology and undertake a second round of path exploration. Consider a node *u*_*z*_ with *p* = *δ* ^−^(*u*_*z*_) incoming nodes, written as 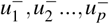, and *q* = *δ* ^+^(*u*_*z*_) outgoing nodes, written as 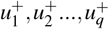. Write 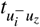 and 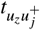 the number of visits to the respective edges connecting *u*_*z*_ in the path, where *i* = 1, …, *p* and *j* = 1, …, *q*. Write 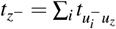 and 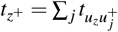 the real indegree and outdegree of the node *u*_*z*_. If the entire graph can be traversed with a Hamiltonian cycle, then for any node *u*_*z*_ in the graph, 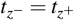,holds. It is important to note the difference between *δ* ^−*/*+^(*u*_*z*_) and 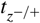,while the former considers only the graph structure, the latter also considers the copy number of nodes. We minimise the objective function: 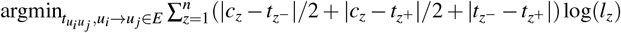. Intuitively, we minimise the sum of the differences between the indegrees and outdegrees, as well as the discrepancies between the expected node copy number and the observed degree. In the objective function, the term for each node is weighted by the logarithm of the corresponding sequence length to assign greater importance to longer sequences. The variables 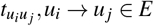 take discrete values and are bounded by 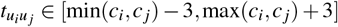. If the solution space is no greater than 100 million, we perform a brute force search; otherwise, we use simulated annealing.

### Assembling plant organelle genomes

We ran Oatk to generate organelle genome assemblies for 195 land plant species sequenced by the Tree of Life programme ^24^. For organellar sequence identification, the Embryophyta plastid and mitochondrial profile HMM gene databases were specified using the ‘-p’ and ‘-m’ parameters, respectively. GenomeScope ^46^ (version 2.0) was employed to estimate the haplotype data coverage of the nuclear genome. The coverage threshold (‘-c’) was set to five times the estimated haplotype coverage, and the number of threads (‘-t’) was set to 6 for running Oatk, with all other parameters left at their default settings.

We ran MBG^25^ (version 1.0.16) and PMAT ^17^ (version 1.5.3) for genome assembly graph construction to compare with the Oatk assembler *syncasm*. For MBG, we used the same *k*-mer size and coverage threshold as *syncasm*, i.e., ‘-k’ 1,001 and ‘-c’ five times the haplotype coverage, and set ‘-t’ to 6 to use six threads. All other parameters were left at their default settings. For PMAT, we configured the genome size parameter (‘-g’) to the estimated value from GenomeScope, set the sequence type parameter (‘-st’) to ‘hifi’, and adjusted the sampling ratio factor parameter (‘-fc’) based on the GenomeScope haplotype coverage estimation to use 3× data. Additionally, we set the ‘-cpu’ parameter to 6 to use six threads and enable the ‘-m’ option to retain sequence data in memory for faster processing. PMAT generates separate assembly graphs for plastid and mitochondrial genomes. However, sequence misclassification often results in a significant proportion of shared sequences between the two assembly graphs. To address this issue, we merged the two graphs to create a unified assembly graph for both organelle genomes, removing redundant sequences and edges while preserving the overall graph structure.

We ran GetOrganelle ^14^ (version 1.7.7.1) to generate organelle genomes from assembly graphs constructed by *syncasm*, MBG, and PMAT. We set ‘-F’ to ‘embplant pt’ and ‘embplant mt’ respectively for assembling plastid and mitochondrial genomes, and set ‘-t’ to 6 to use six threads. We ran the Oatk *hmmannot*-*pathfinder* pipeline to compare with GetOrganelle. This was achieved by using the ‘-G’ parameter in Oatk, which allows it to take an existing assembly graph as input. By specifying this parameter, Oatk bypasses the assembly graph construction from the raw sequence data.

### Calculating SC and IR component sizes for plastomes

We ran NUCmer ^47^ (version 4.0.0rc1) with the parameter ‘-- maxmatch’ to generate a self-alignment for the genome sequence. Alignments with sequence identity below 95% and those not in the reverse-complement orientation were filtered out. Alignments within a distance smaller than 50 bp were merged, where the distance between two alignments is defined as the Manhattan distance between their closest endpoints. Following merging, the two largest aligned segments are identified as IRs. The regions flanking these IRs are identified as LSC and SSC based on their respective sizes. The genome sequence is finally rotated so that it starts at the beginning of the *psbA* gene, which is at the start of the LSC, and ends with an IR.

### Mapping reads to assembly graphs

We ran GraphAligner ^48^ (version 1.0.19) to map reads to the assembly graphs with the parameters ‘--precise-clipping 0.9 --min-alignment-score 5000 -x vg’. Alignments were filtered out if less than 95% of the read was aligned to a unique position. GraphAligner outputs a path on the graph for each alignment. For any given graph path of interest, we examine the alignment path of each read and count it as a supporting read if the alignment path fully covers the target graph path. Alignments in both directions are considered, and the sum is taken as the total number of supporting reads.

### Identifying shared sequences between plastid and mitochondrial genomes

We ran NUCmer ^47^ (version 4.0.0rc1) with the parameter ‘--maxmatch’ to map the plastome to the mitogenome. We then ran *delta-filter* ^47^ (version 4.0.0rc1) with the ‘-l 1000’ parameter to remove short alignments, the ‘-i 90’ parameter to retain only alignments of at least 90% sequence identity, and the ‘-r’ parameter to keep only the best mitogenome match to each region of plastome sequence. The sequences corresponding to the resulting alignments are considered the shared sequences between the two organelle genomes.

### Constructing the phylogenetic tree for the *psaA* gene from shared gene sequences

We ran *hmmannot* to generate gene annotations of the shared sequences and extracted sequences annotated as *psaA* genes. We ran MAFFT ^43^ (version 7.505) with the parameters ‘--localpair --maxiterate 1000’ to generate a multiple alignment of these gene sequences and ran IQ-TREE ^49^ (version 2.3.4) with default parameters to generate a phylogenetic tree. The calculations of distances between tree nodes and identification of internal MRCA nodes were performed using Bio.Phylo functions ^50^ in Biopython (version 1.83).

## Data availability

The source code for Oatk is available in the GitHub repository https://github.com/c-zhou/oatk. The source code for OatkDB and the profile HMM gene database are available in the GitHub repository https://github.com/c-zhou/OatkDB. Raw sequence data are available in public data repositories, with accession numbers listed in the Supplementary Table S3. The organelle genome assembly data for all species are available in the Zenodo repository https://doi.org/10.5281/zenodo.13952353

## Authors’ contributions

All authors contributed ideas. CZ implemented Oatk and obtained the assembly results. MBr provided initial computational solutions to some aspects. CZ drafted the paper with RD. All authors commented on and approved the final manuscript.

## Acknowledgements

We thank Sergey Nurk and Alex Twyford for discussions, Yaqi Su for exploration and feedback on early Oatk results, Ksenia Krasheninnikova for the Nextflow implementation, and Lia Obinu for the Bioconda implementation. For the purpose of open access, the author has applied a CC BY public copyright licence to any Author Accepted Manuscript version arising from this submission.

