## supplementary_figures for "Oatk: a de novo assembly tool for complex plant organelle genomes"

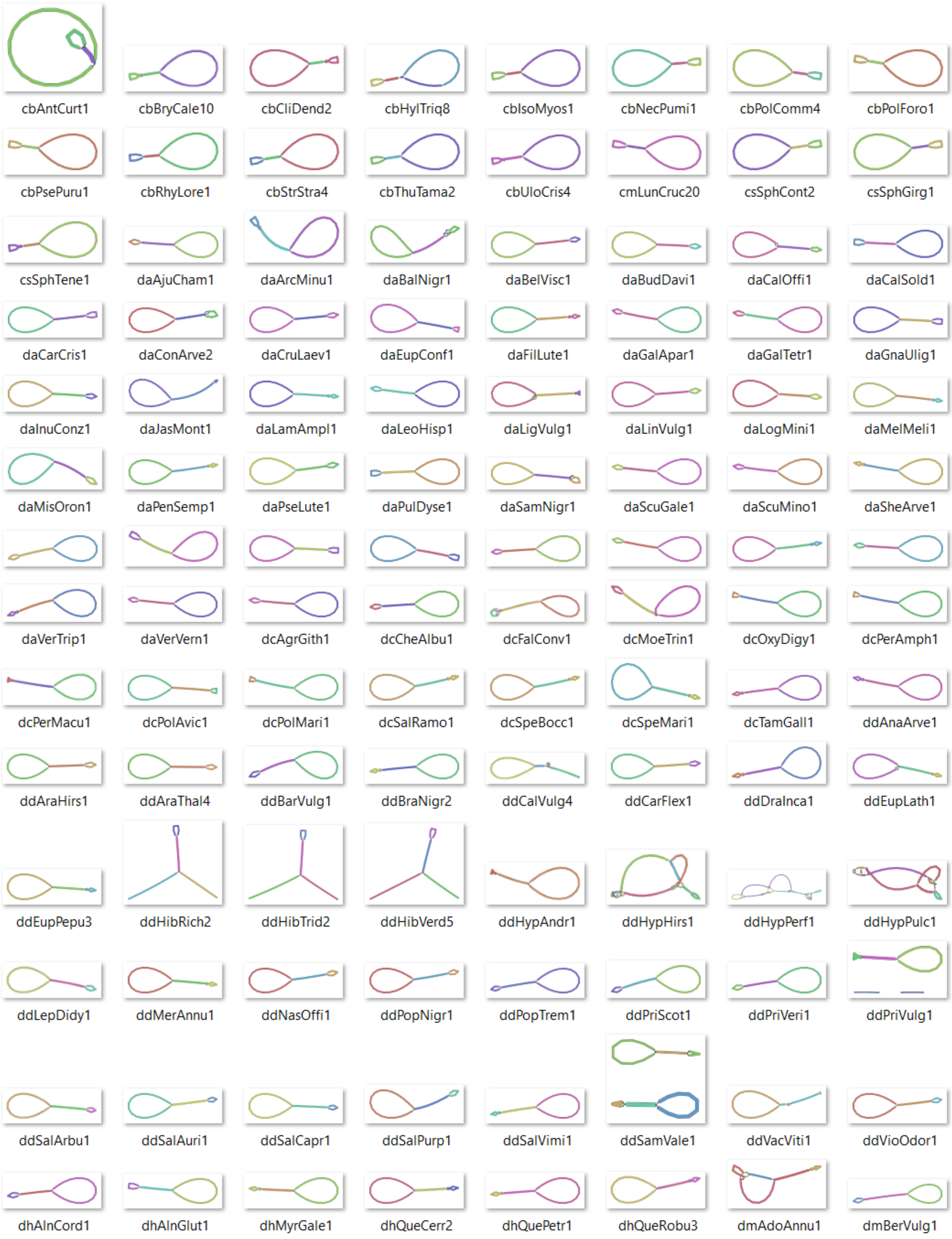


**Figure S1.** continued on next page

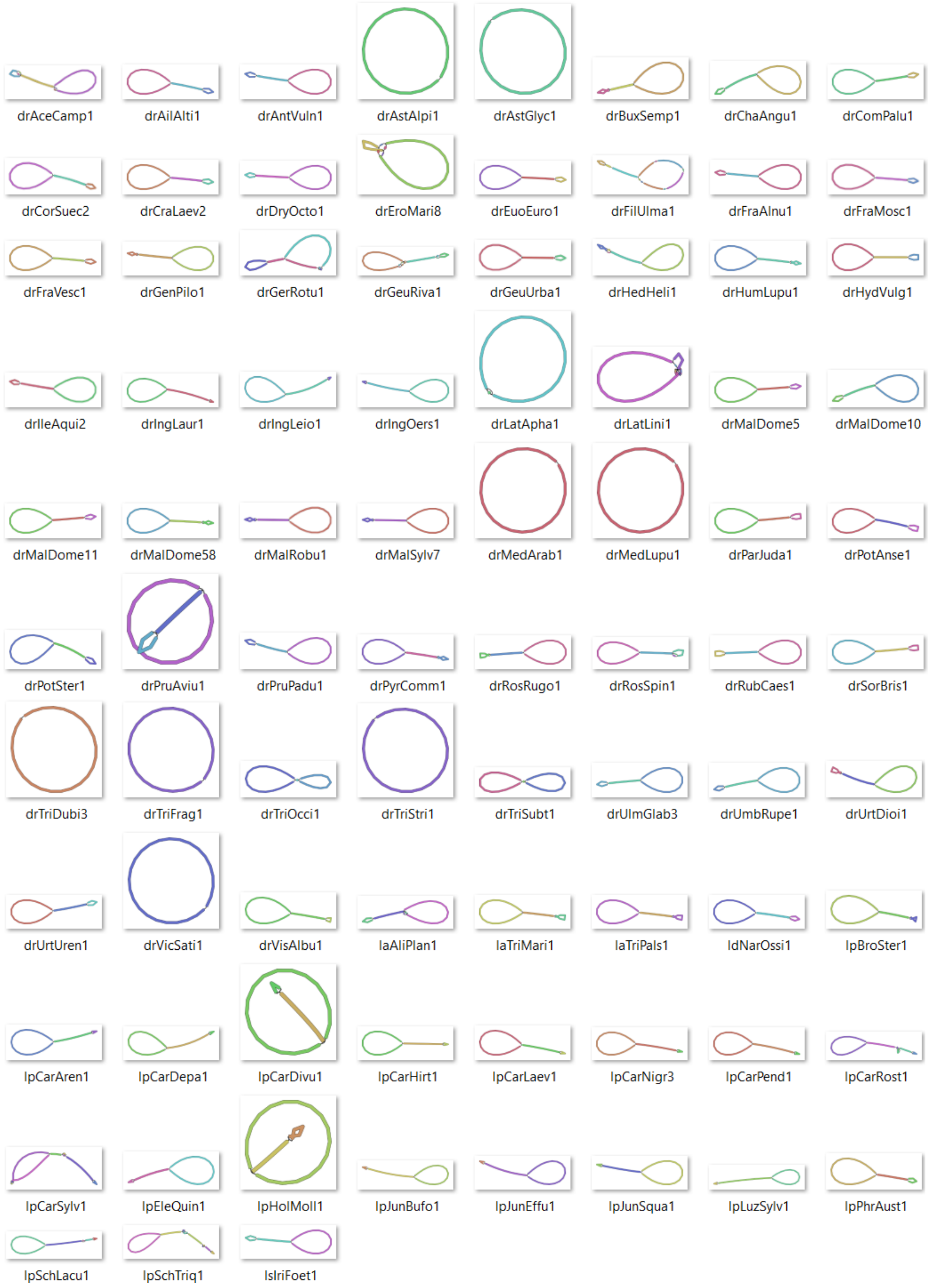


**Figure S1. Assembly graphs for all plastomes.** The graphs were produced using Bandage[1].

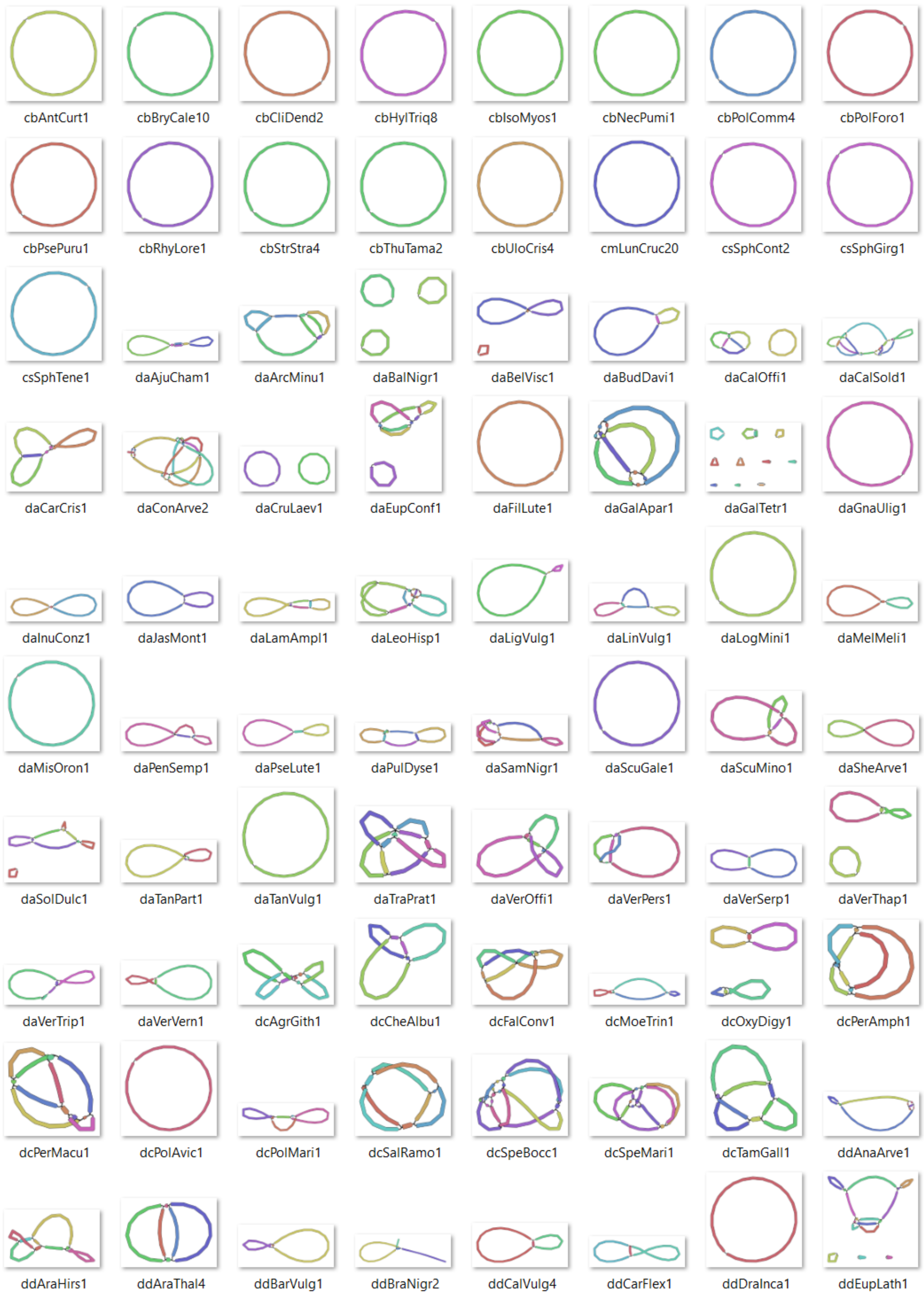


**Figure S2.** continued on next page

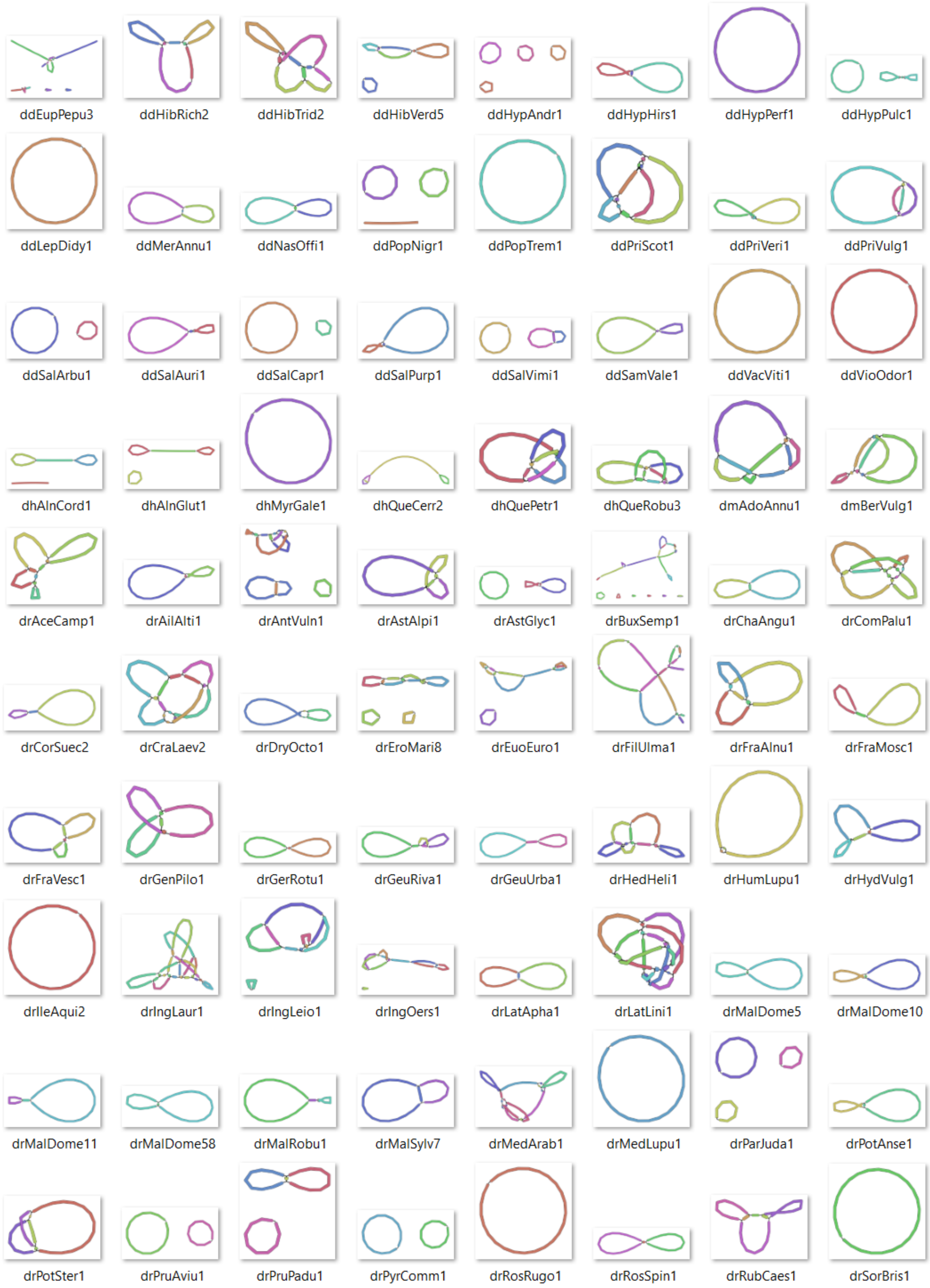


**Figure S2.** continued on next page

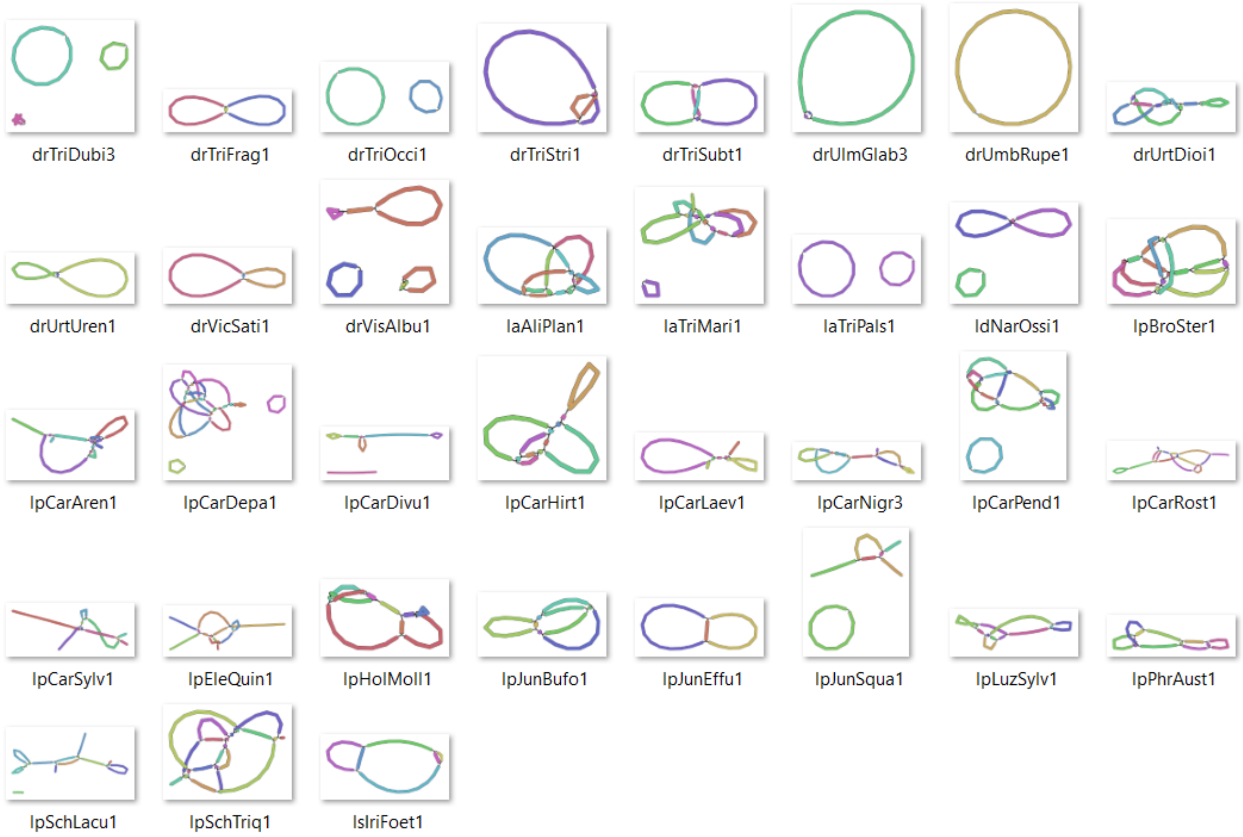


**Figure S2. Assembly graphs for all mitogenomes.** The graphs were produced using Bandage[1].

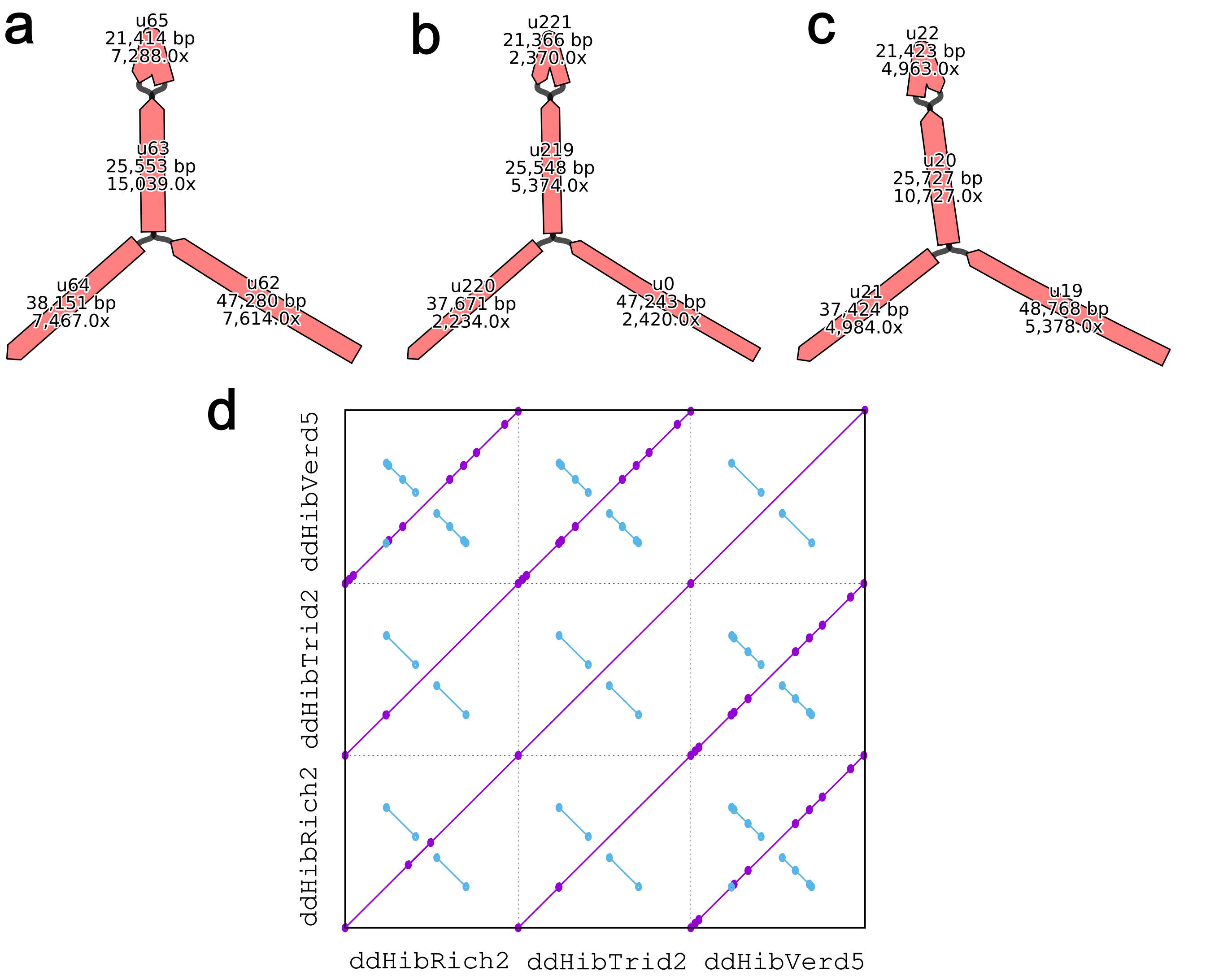


**Figure S3. Plastome assembly results for three hibiscuses. a-c**, the genome assembly graph for *Hibiscus richardsonii* (**a**), *Hibiscus tridactylites* (**b**) and *Hibiscus verdcourtii* (**c**). **d**, a collinearity plot between the three sequences. The assembly graphs were produced using Bandage[1] with additional manual adjustments.

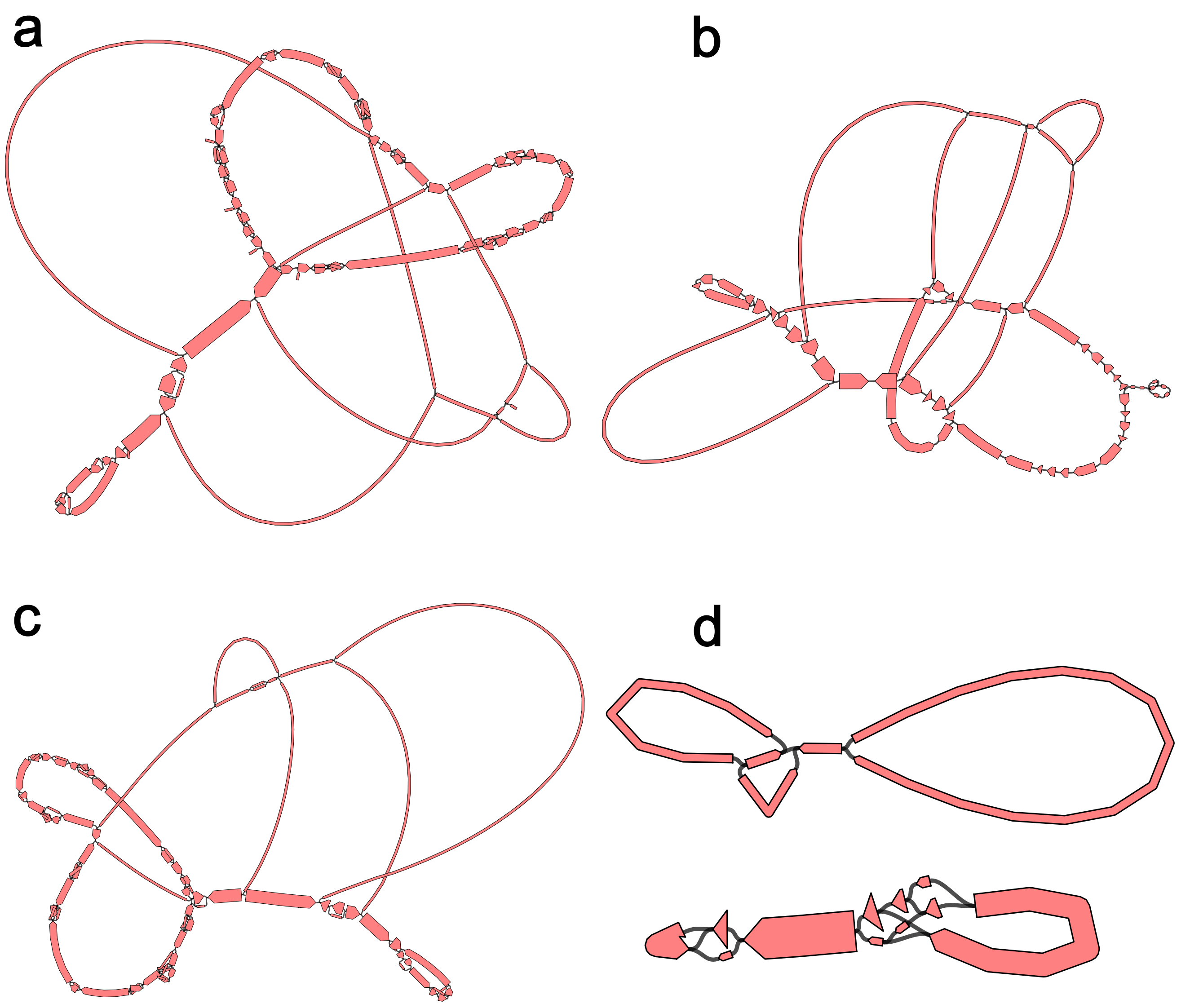


**Figure S4. Example mixture assembly graphs generated by different assemblers for *Geum rivale*. a**, MBG graph. **b** PMAT graph. **c**, Syncasm raw graph. **d**, Syncasm disentangled graph. In each graph, the width of the bar is proportional to the sequence coverage. The coverage of plastome sequences is higher than that of the mitogenome sequences. The assembly graphs in **a-c** represent mixtures of plastid and mitochondrial genomes. Syncasm successfully separates the two organelle genomes after disentangling (**d**). The assembly graphs were produced using Bandage[1] with additional manual adjustments.

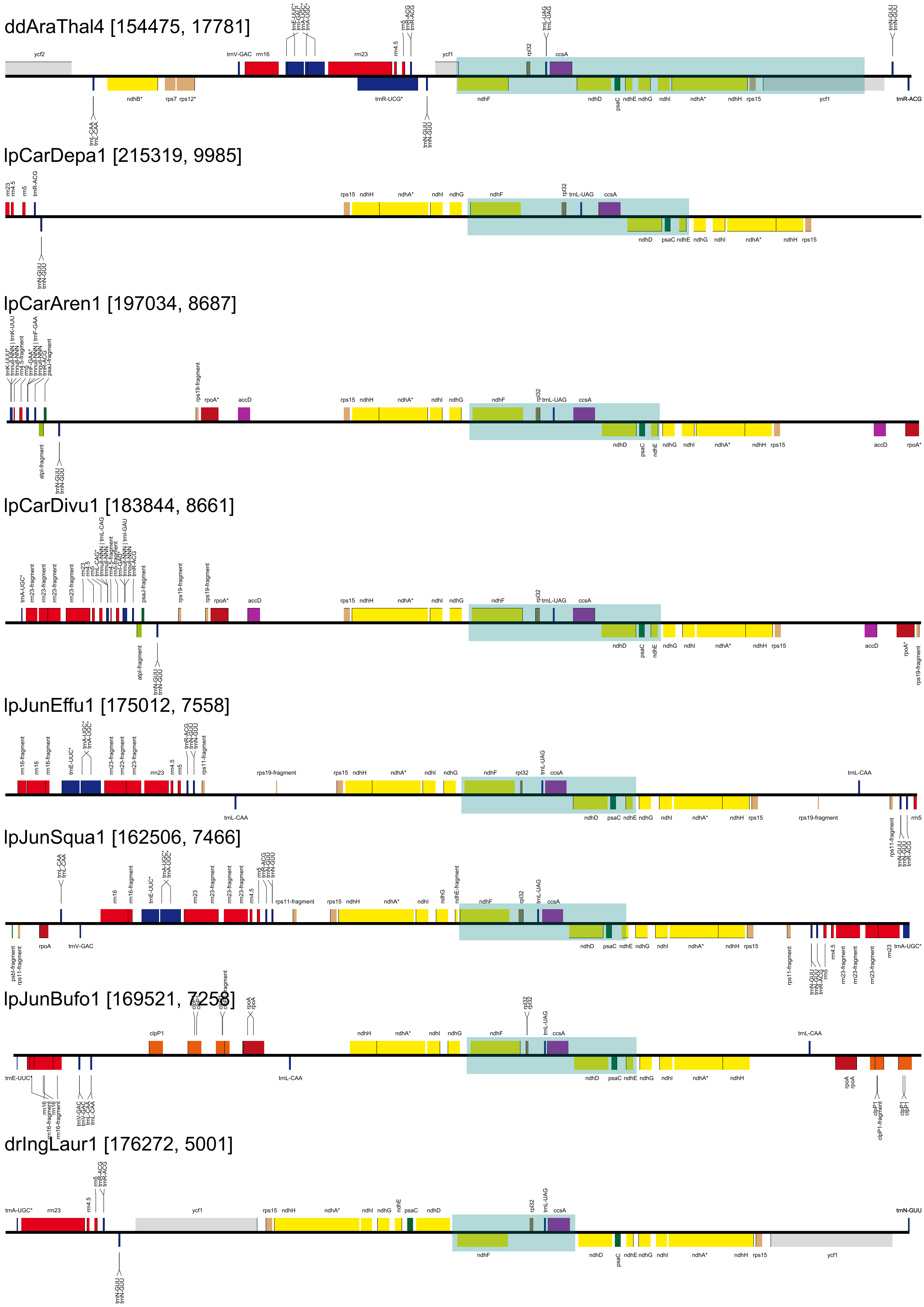


**Figure S5.** continued on next page

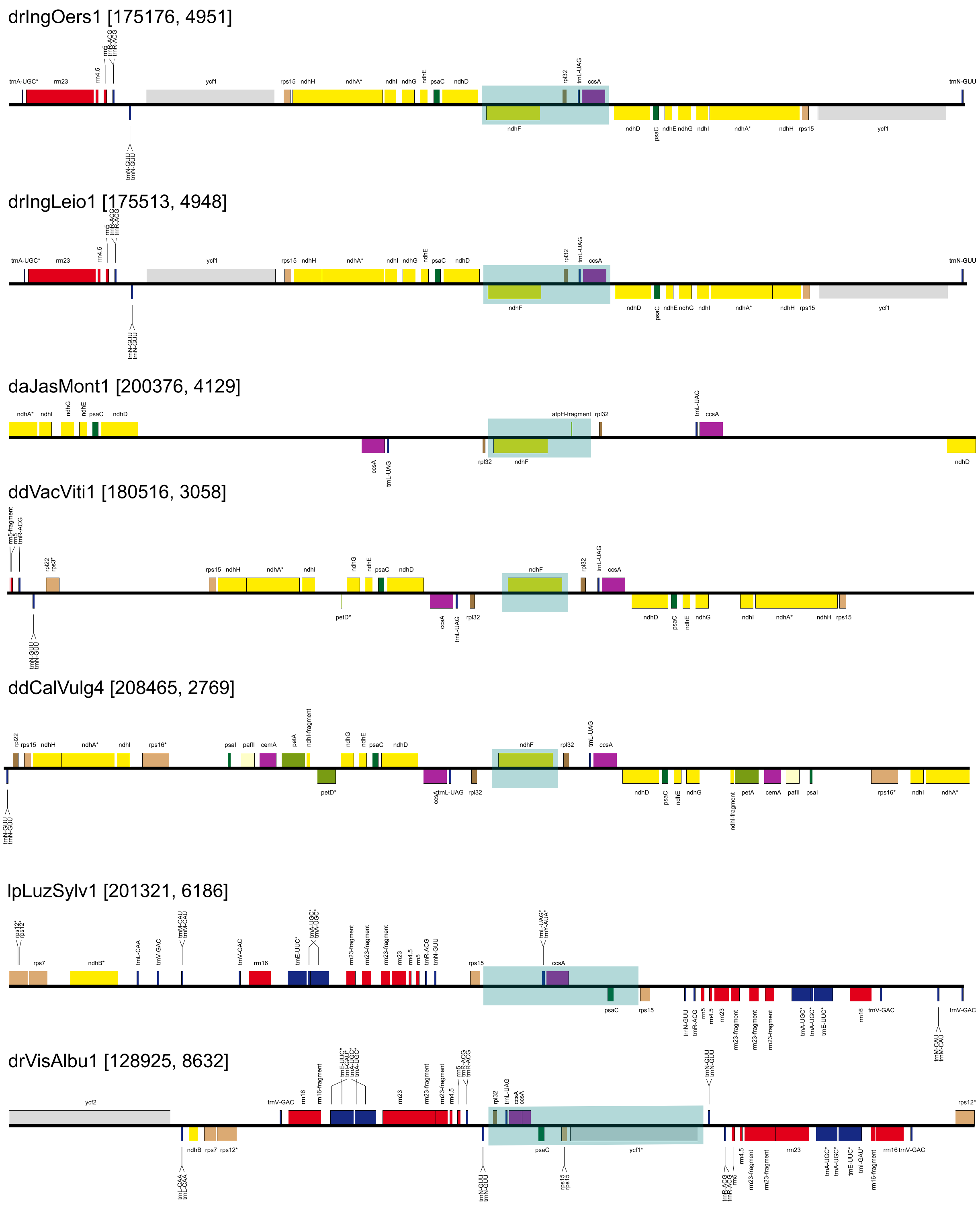


**Figure S5. Gene annotation results in the SSC region for species with SSCs smaller than 10 kb.** The numbers in square brackets next to the species ID denote the genome size and the SSC size. The light blue box represents the approximate location of the SSC. The annotation was performed using GeSeq[2] and visualised with OGDRAW[3].

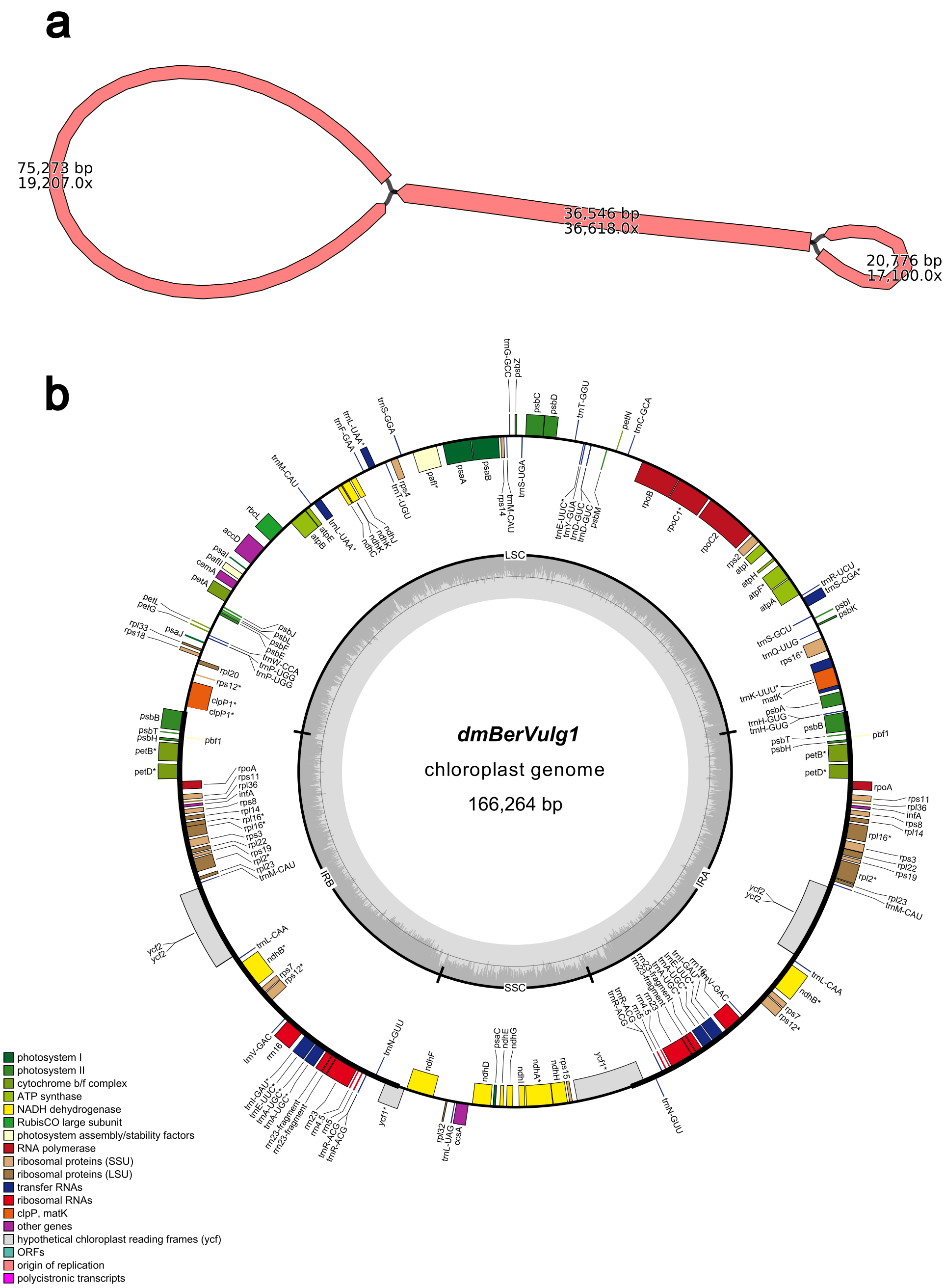


**Figure S6.** **Plastome structure of *Berberis vulgaris*.** **a**, the genome assembly graph. The numbers on the bar indicate the sequence length and coverage. **b**, the genome annotation. The assembly graph was produced using Bandage[1] with additional manual adjustments. The annotation was performed using GeSeq[2] and visualised with OGDRAW[3].

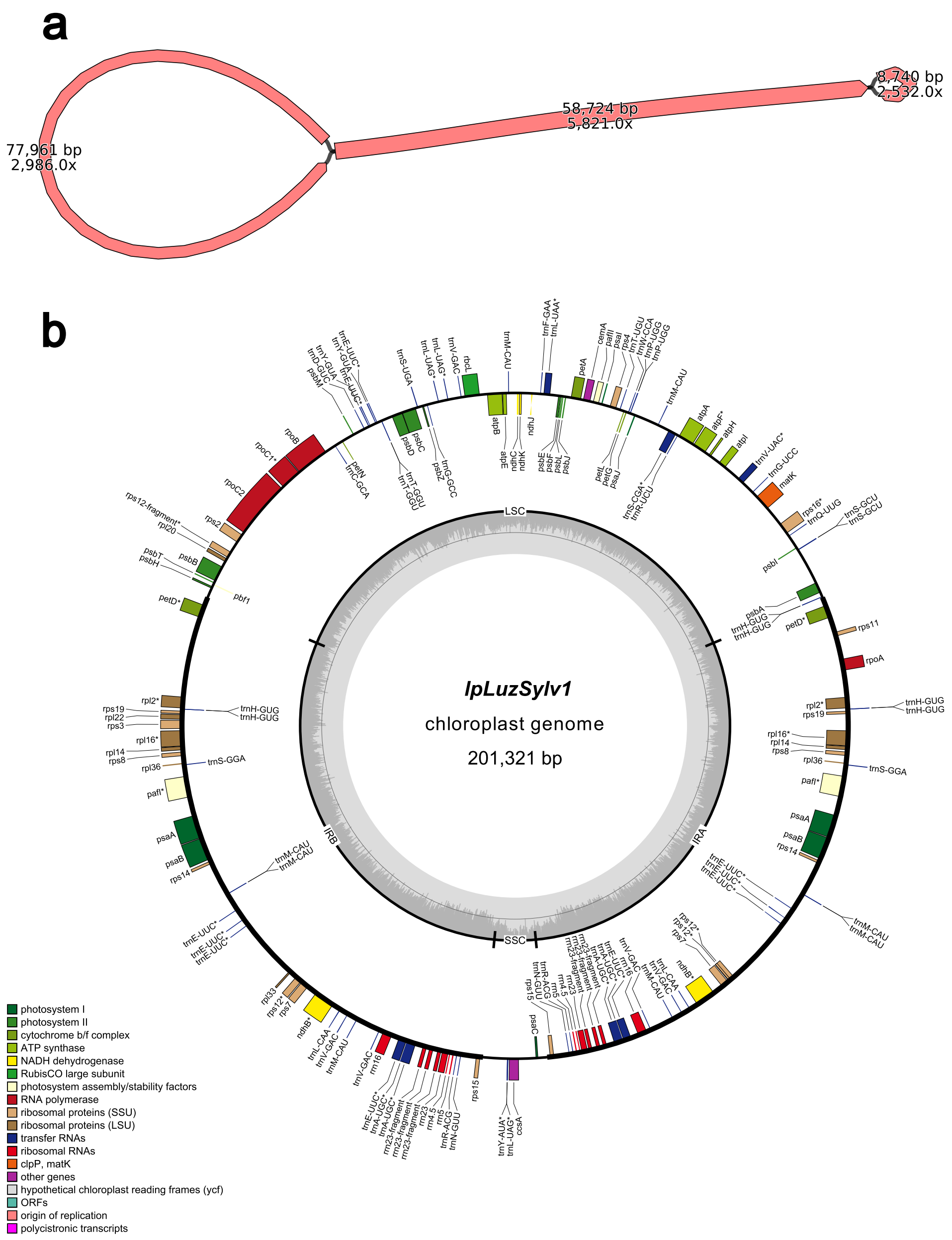


**Figure S7. Plastome structure of *Luzula sylvatica*. a**, the genome assembly graph. The numbers on the bar indicate the sequence length and coverage. **b**, the genome annotation. The assembly graph was produced using Bandage[1] with additional manual adjustments. The annotation was performed using GeSeq[2] and visualised with OGDRAW[3].

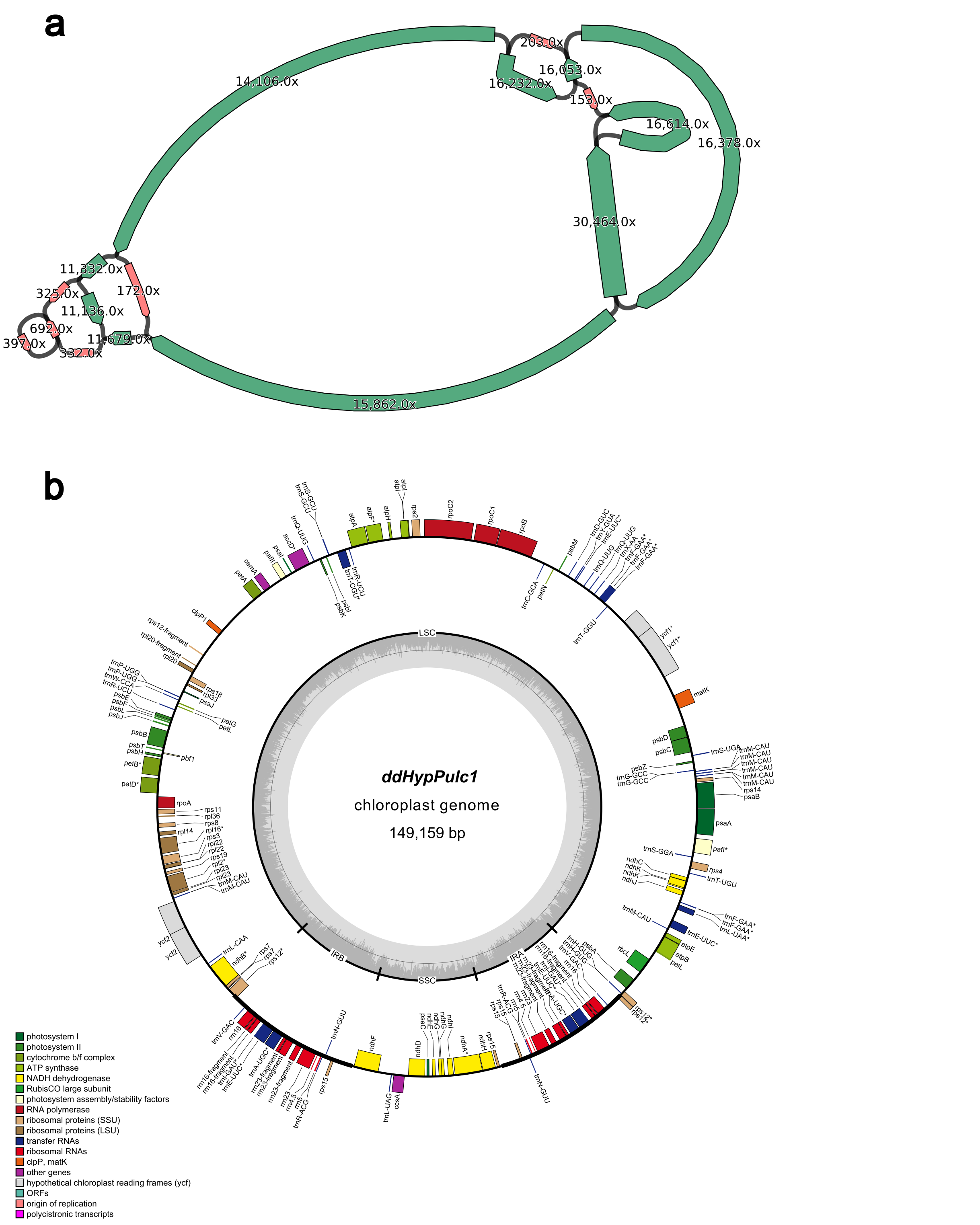


**Figure S8. Plastome structure of *Hypericum pulchrum*. a**, the genome assembly graph. The numbers on the bar indicate the sequence length and coverage. Sequences in green represent a path in the graph for the most abundant haplotype, while sequences in red represent heteroplasmy. **b**, the genome annotation. The assembly graph was produced using Bandage[1] with additional manual adjustments. The annotation was performed using GeSeq[2] and visualised with OGDRAW[3].

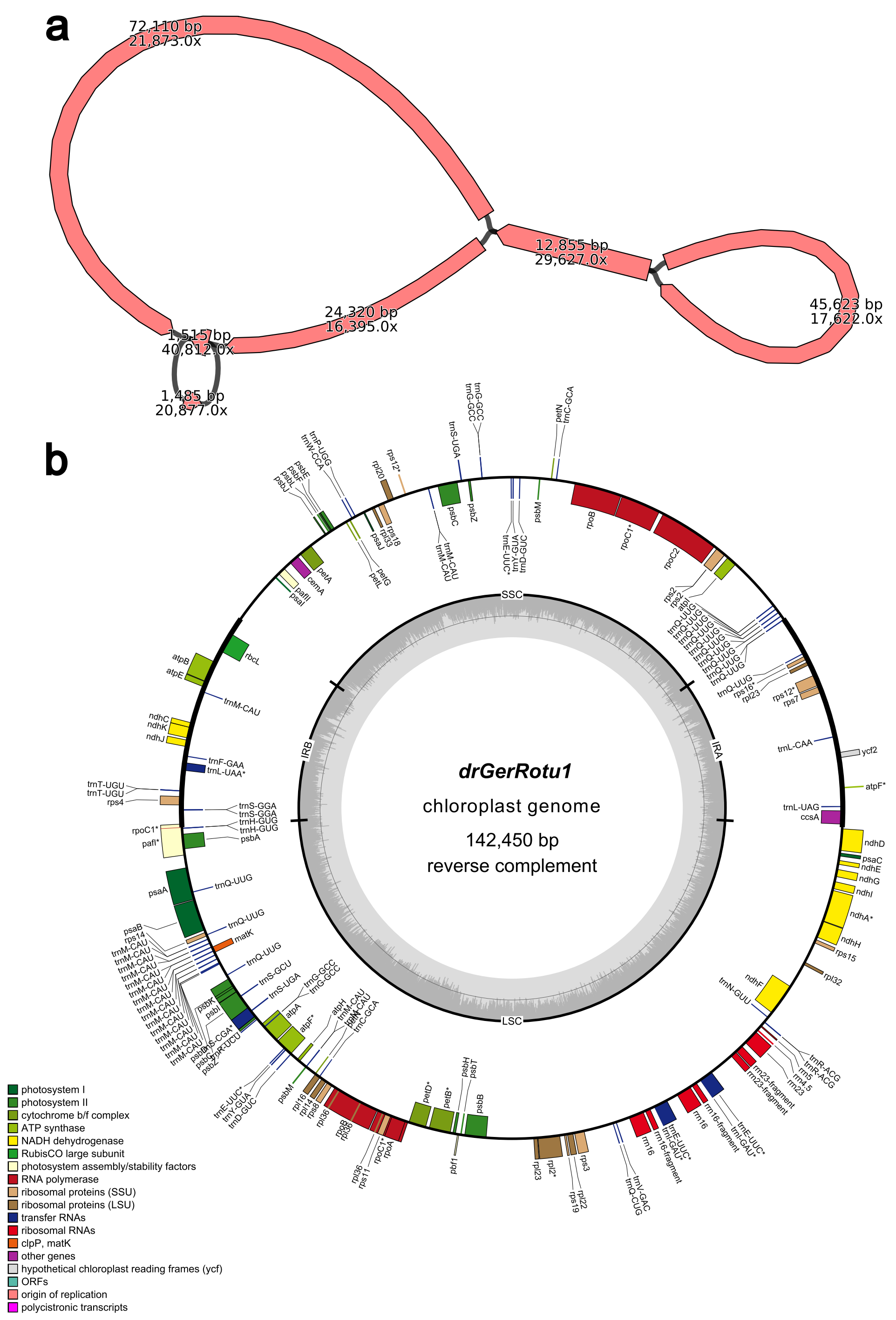


**Figure S9. Plastome structure of *Geranium rotundifolium*. a**, the genome assembly graph. The numbers on the bar indicate the sequence length and coverage. **b**, the genome annotation. The assembly graph was produced using Bandage[1] with additional manual adjustments. The annotation was performed using GeSeq[2] and visualised with OGDRAW[3].

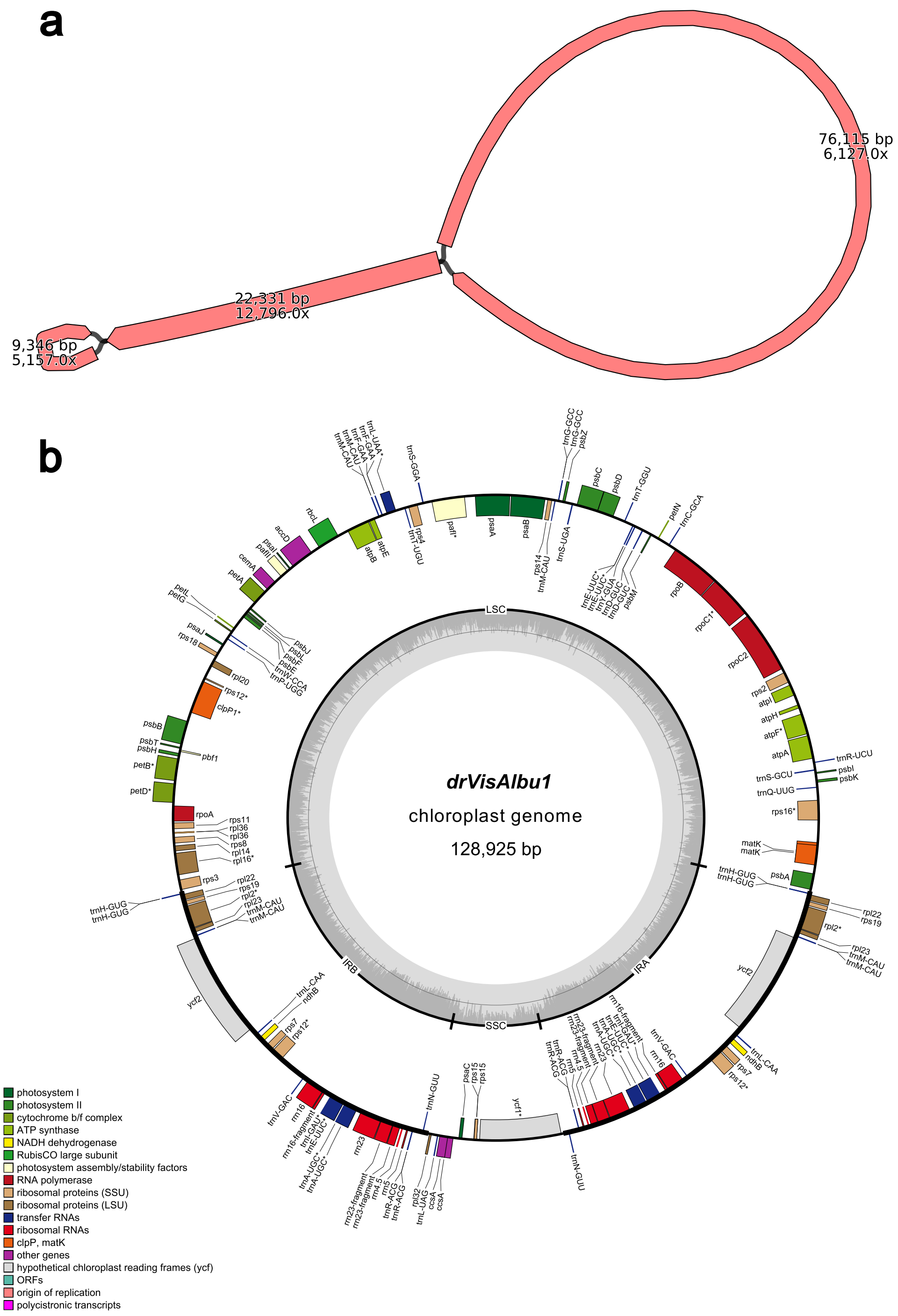


**Figure S10. Plastome structure of *Viscum album*. a**, the genome assembly graph. The numbers on the bar indicate the sequence length and coverage. **b**, the genome annotation. The assembly graph was produced using Bandage[1] with additional manual adjustments. The annotation was performed using GeSeq[2] and visualised with OGDRAW[3].

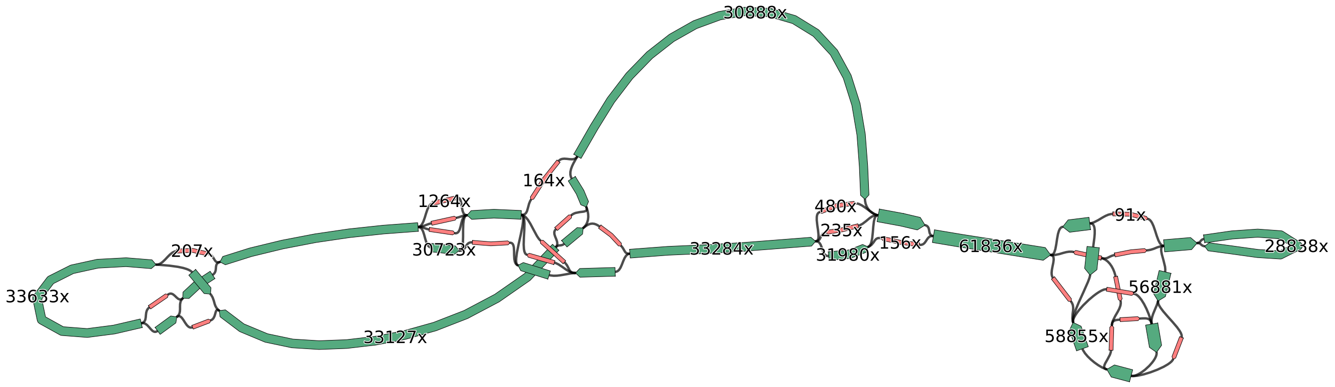


**Figure S11. Plastome structure of *Hypericum perforatum*.** In the graph, the width of each bar is proportional to the sequence coverage. The number on the bar indicates the actual coverage of the sequence, with some omitted for clarity. Sequences in green represent a path in the graph for the most abundant haplotype, while sequences in red represent heteroplasmy. The assembly graph was produced using Bandage[1] with additional manual adjustments.

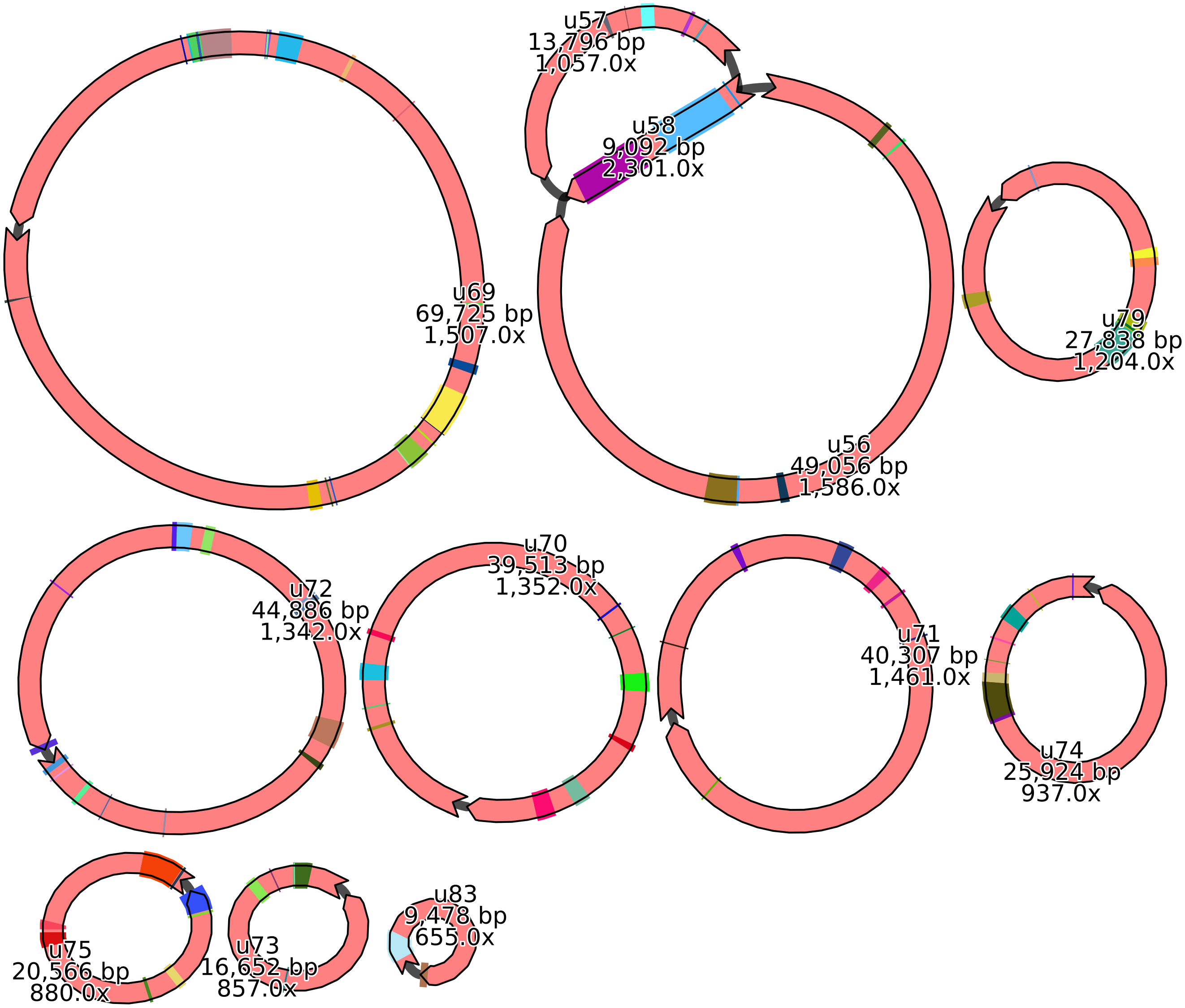


**Figure S12. Mitogenome assembly of *Galeopsis tetrahit*.** The genome assembly consists of ten components, with coloured blocks over the bars representing annotated genes. The numbers on the bar indicate the sequence name, length and coverage. The assembly graph was produced using Bandage[1] with additional manual adjustments.

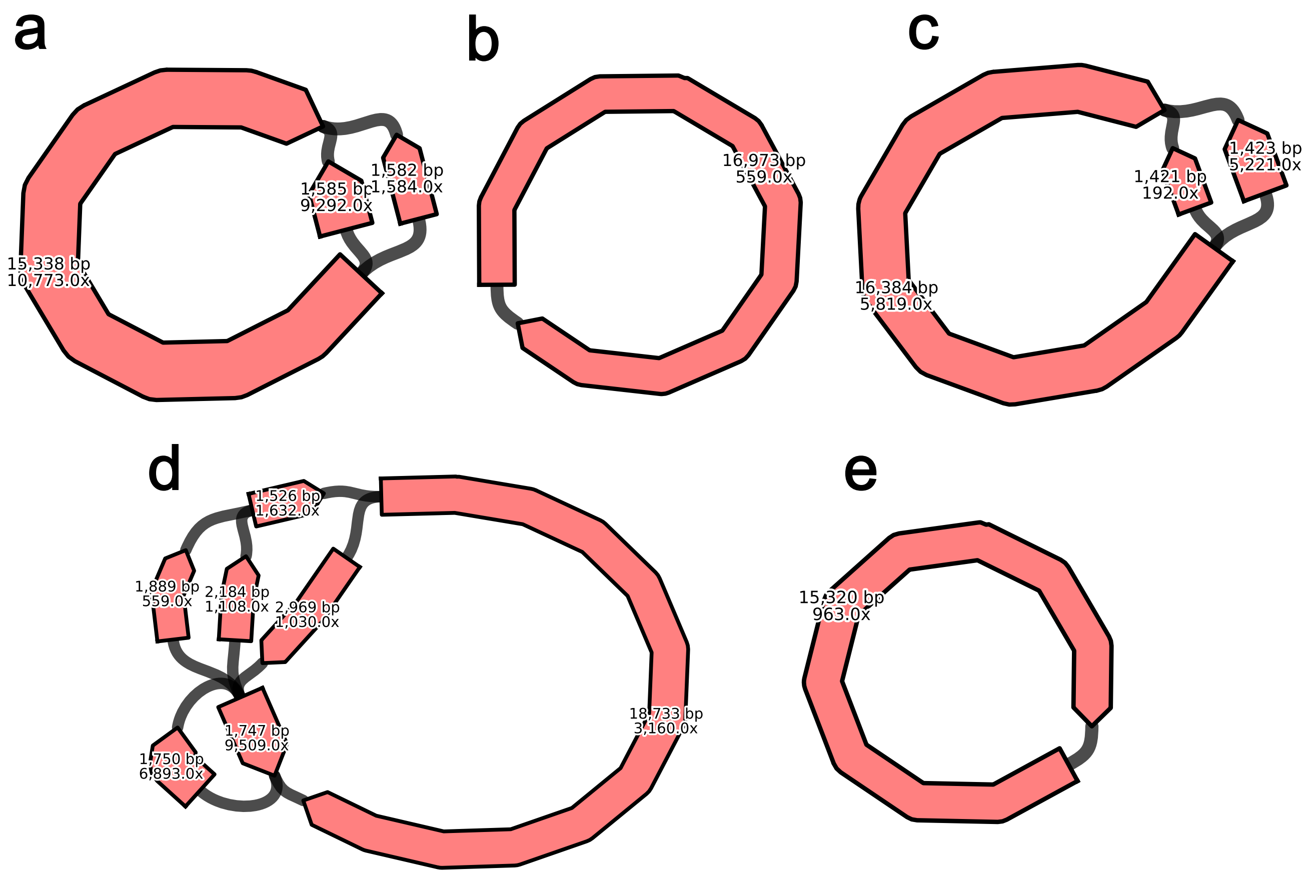


**Figure S13. Mitogenome assembly of five animal species. a**, *Micromys minutus* (TOLID: mMicMin1; ACCESSION: PRJEB72093). **b**, *Abramis brama* (TOLID: fAbrBra2; ACCESSION: PRJEB73975). **c**, *Anas platyrhynchos* (TOLID: bAnaPla2; ACCESSION: PRJEB76742). **d**, *Podarcis gaigeae* (TOLID: rPodGai1; ACCESSION: PRJEB76108). **e**, *Erebia medusa* (TOLID: ilEreMedu1; ACCESSION: PRJEB76717). The numbers on the bar indicate the sequence length and coverage. The assembly graph was produced using Bandage[1] with additional manual adjustments.
